# Bivalent ligands as a new universal chemotype of BCL6 degrader by inducing aggregation

**DOI:** 10.1101/2025.02.25.638777

**Authors:** Rong Hu, Xia-Tong Hu, Ying-Yue Yang, Peng Luo, Cai-Hua Li, Zhi-Hao Liu, Luo-Ting Yu, Ning-Yu Wang

## Abstract

BCL6 naturally exists as a homodimer to exert its transcriptional repressive function. Previous studies shown that molecular glue BI3802 can induce BCL6 dimer polymerization and thus degradation by ubiquitin-proteasome system. In this study we proposed a new strategy to degrade BCL6 by its homo-bivalent ligands, which can be rationally designed by assemble two BCL6 BTB ligands via linker. These homo-bivalent degraders (HBiDs) can induce polymerization of BCL6 dimer through chain assembly and ultimately lead to its ubiquitination and degradation. Since HBiD does not require a fragment to interact with components of protein degradation system, it could exhibit better target selectivity than traditional protein degraders while avoid drug resistance caused by mutations in protein degradation components. Here we will provide the proof-of-concept evidences for HBiD as BCL6 degrader. Most HBiDs synthesized in this study could degrade BCL6 regardless of the chemotype of BCL6 ligands, with extraordinary target selectivity. The degradation of BCL6 depends on the aggregation of BCL6 dimer induced by HBiD and the recruitment of SIAH1, the natural E3 ligase of BCL6. As protein dimerization is a common phenomenon in cells, HBiD may provide us with a powerful strategy for designing protein degraders as drug candidates or molecular probes.

## Introduction

Small molecule protein degraders can downregulate the abundance of protein of interesting (POI) with the help of human protein balance systems, such as the ubiquitin-proteasome system (UPS) or the Autophagic Lysosomal System (ALS)^[1-2]^. At present, many types of protein degraders have been reported, including proteolysis-targeting chimeras (PROTAC)^[3-4]^, hydrophobic tags (HyTs)^[5]^, molecular glues^[6]^, lysosome-targeting chimaeras (LYTAC)^[7-8]^, autophagy-targeting chimera (AUTAC)^[9-10]^, autophagosome-tethering compound (ATTEC)^[11]^, and AUTOphagy-TArgeting Chimera (AUTOTAC)^[12]^. Although hundreds of pathogenic proteins have been targeted by these protein degradation technologies, the vast majority of proteins in the human proteome have not yet been targeted for degradation by small molecules^[13-14]^. The development of protein degraders with new mechanisms or modes of action can not only enable more proteins to be targeted for degradation, but also partially solve the problems faced by current protein degradation technologies, such as poor target selectivity and drug resistance^[15-16]^.

B cell lymphoma 6 (BCL6) is a proto-oncogene found in adult and childhood cancers and was first discovered in diffuse large B-cell lymphoma (DLBCL)^[17]^, its exact physiological function in human cells has not been fully elucidated. The structure of BCL6 is mainly divided into three parts^[18]^: (1) a C-terminus C_2_H_2_-type zinc finger domain which can identify specific DNA sequences; (2) the central transcriptional repressor (RD2) domain associated with the activity and stability of BCL6; and (3) the N-terminus Broad-Complex, Tramtrack and Bric a brac/poxvirus and zinc finger (BTB/POZ) domains for recruitment of transcriptional co-repressor complex. BCL6 usually performs its physiological function as dimer, the BTB domains on the two protein subunits form a groove at the junction of the protein subunits through a tight winding from head to tail, facilitating the protein-protein interactions of BCL6 with transcriptional corepressors such as NCOR1, SMRT, and BCOR, which result in the transcriptional inhibition of tumor suppressor genes, high fidelity DNA repair genes, and cell differentiation related genes^[19]^.

Recent studies reported that the small molecule BI3802 can degrade BCL6 as a molecular glue. It could induce higher order assembly of BCL6 dimers in a face-to-face binding mode to form filaments through hydrophobic interactions. These BCL6 filaments could be identified by SIAH1, an endogenous E3 ubiquitin ligase of BCL6, and induced proteasome degradation^[20]^.

Since both BTB domains in a BCL6 dimer could bind a ligand, we hypothesize that a bivalent BCL6 ligand, which contains two BCL6 ligand fragments, might bind to two BCL6 dimers and also induce the polymerization of BCL6 dimer through chain assembly and finally induce the degradation of BCL6 by UPS (Figure 1A). This kind of HBiDs only relies on the affinity between the ligand fragment and the BCL6 BTB domain, rather than the key hydrophobic interactions on which molecular glues depend. Theoretically, any reported BCL6 BTB inhibitor is expected to be readily transformed into HBiDs, which will greatly expand the chemical structure space of current BCL6 degraders. In addition, HBiDs are expected to exhibit extremely high target selectivity due to its mode of action depends on the presence of the target protein dimer, the appropriate ligand-to-target affinity, the appropriate length and rigidity of the linker as well as the moderate stability of the target protein multimer.

**Figure 1.**
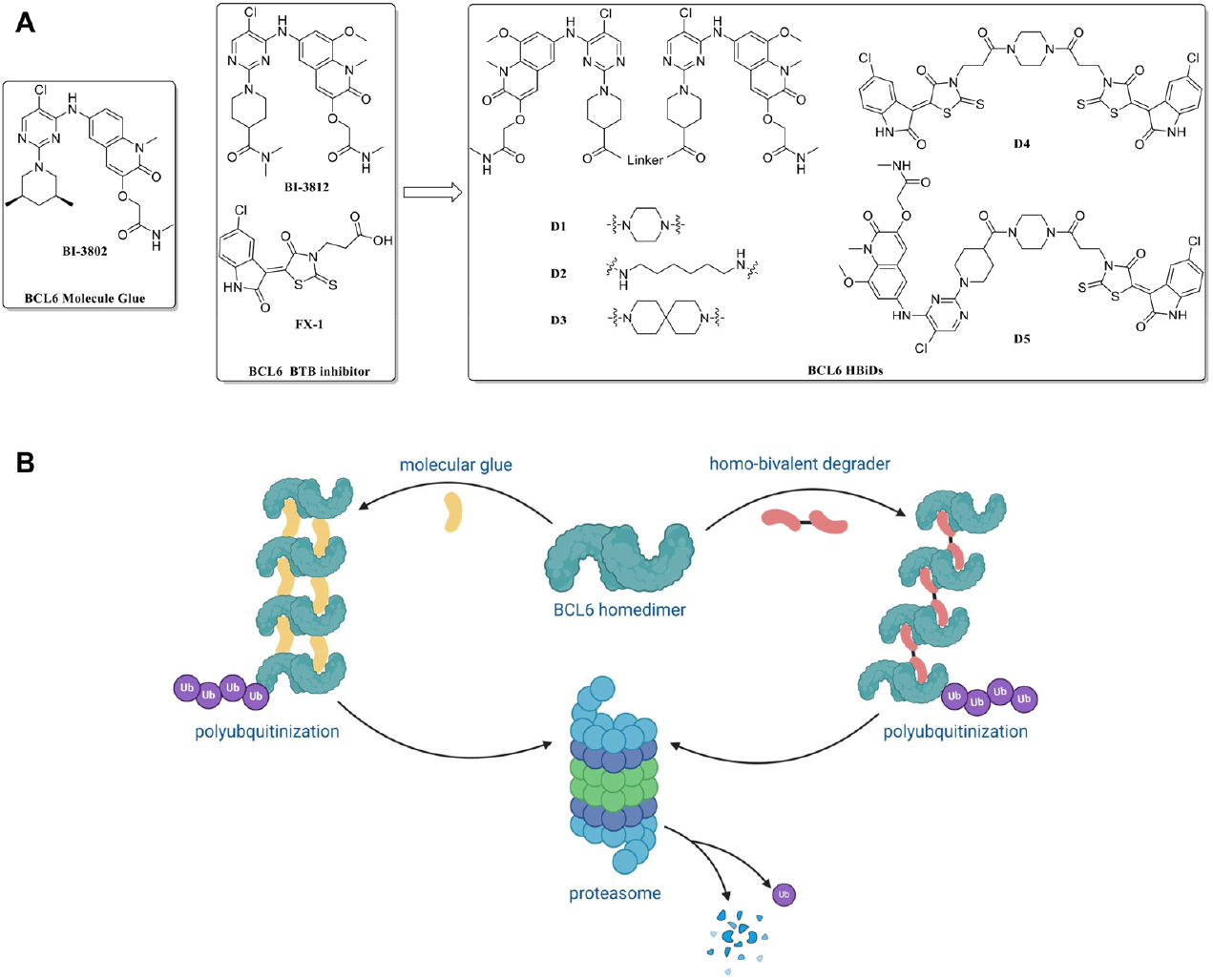
Discovery of HBiD. A. The design of BCL6 HBiDs. B. Schematic presentation of the mechanism of BCL6 degradation by molecular glue or HBiD.

## Results

HBiDs (**D1**-**D4**) based on two structurally unrelated BCL6 inhibitors BI3812 and FX-1 as well as the hetero-bivalent ligand **D5** were synthesized in this study (Figure 1B). We firstly evaluated the degradation ability of all HBiDs in Farage cells, a DLBCL cell line expressing a high level of BCL6^[21]^. As expected, most HBiDs with a rigid linker could degrade BCL6 efficiently regardless of the structural type of BCL6 ligands, while **D2** with a flexible linker failed to degrade BCL6 even at a concentration of 10 μM, which might be due to the adverse effect of flexible linker on the stability of protein polymers. The inability of the hetero-bivalent ligand **D5** to degrade BCL6 might be due to the difference in target affinity between these two BCL6 ligand fragments, which result in the unstable assembly of BCL6 dimer into polymers (Figure 2A). We further evaluated the degradation ability of **D1**/**D3** in three DLBCL cell lines (Farage, SUDHL4, OCI-LY1) in a wider range of concentrations. **D1**/**D3** could degrade BCL6 in a concentration-dependent manner in these cell lines and exhibited a significant hook effect, a typical feature of chimeric molecules such as PROTAC, suggested that HBiDs did function as a bifunctional connector of two BCL6 BTB domains but not a molecular glue to exert its protein degradation activity (Figure 2B/S1A). The HBiD **D4** derived from a weaker BCL6 inhibitor FX-1 also exhibited a weaker degradation ability, suggesting that the affinity of the ligand to the target protein will also significantly affect the protein degradation ability of HBiD (Figure S1B). Since HBiDs do not contain a metabolically unstable hydrophobic fragment that is essential for BCL6 molecular glues like BI3802, we speculate that HBiDs can exhibit better catalytic efficiency and duration to degrade BCL6, and display better protein degradation activity as a result. As expected, **D3** can effectively induce BCL6 degradation than BI3802, with DC_50_ of 19.42 pM in Farage cells after drug exposure for 24 h (Figure S1C). Moreover, the drug exposure for 0.5 h was enough to cause the down-regulation of BCL6, and the protein degradation could last for more than 72 hours (Figure 2B).

**Figure 2.**
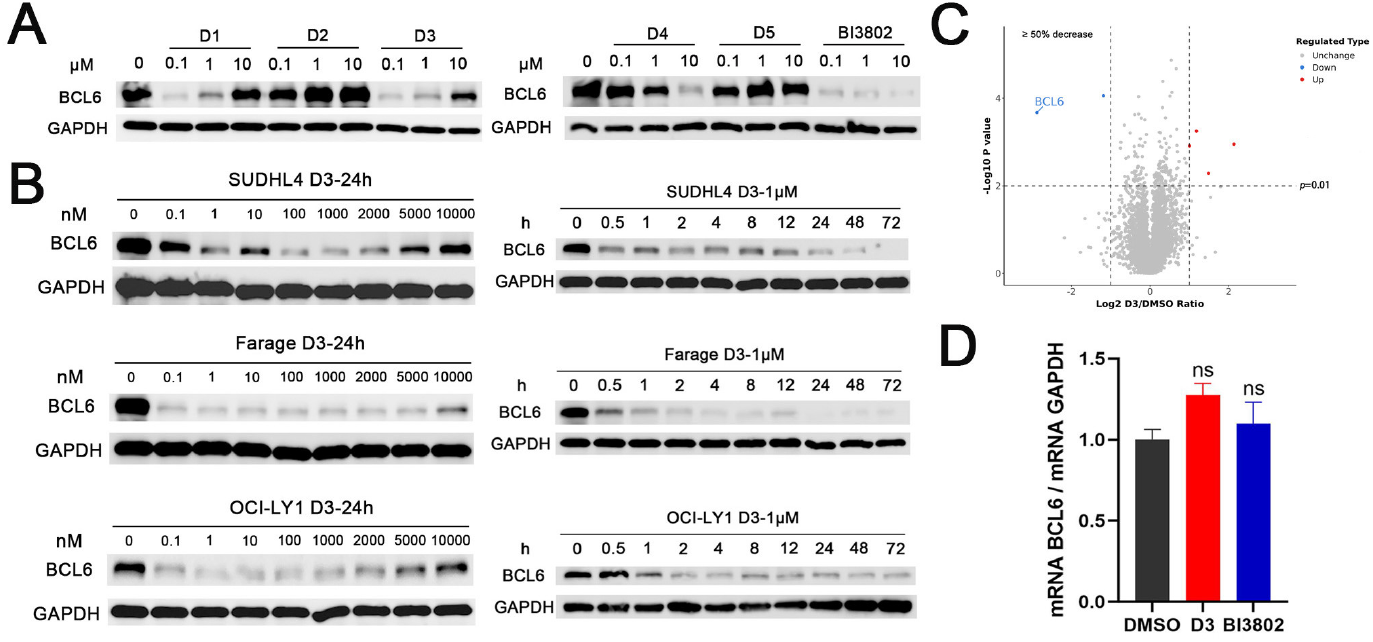
HBiDs efficiently degrade BCL6. A. Immunoblots of BCL6 in Farage cells after treatment with different concentrations of HbiDs for 24 h. B. Immunoblots of BCL6 in Farage, SUDHL4 and OCI-LY1 cells after treatment with different concentrations or times of **D3**. C. Whole-proteome quantification of Farage cells treated with 1 μM **D3** (n = 3) or DMSO (n = 3) for 24 hours (two sided moderated t-test, n = 3). D. mRNA levels of BCL6 quantified by qPCR in Farage cells after treatment with 1 μM **D3**/BI3802 or DMSO for 24 hours. ^n.s.^P > 0.05.

To determine the specificity of **D3** as a BCL6 degrader, we performed quantitative mass spectrometry (MS) based proteomics in Farage cells following **D3** treatment for 24 hours. **D3** could selectively catalyzes degradation of BCL6 over other proteins (Figure 2C). Quantitative real-time PCR showed that both **D3** and BI3802 did not affect the mRNA level of BCL6 (Figure 2D), indicating that the down-regulation of BCL6 was not caused by its transcriptional inhibition.

**D3**-induced BCL6 degradation could be restored by the 26S proteasome inhibitor MG132 or the ubiquitin activating enzyme UBA1 inhibitor MLN7243 in Farage cells, but not by autophagy inhibitor chloroquine or the neddylation pathway inhibitor MLN4924, suggesting that this degradation is dependent on the UPS but not mediated by the Cullin-RING E3 ubiquitin ligases, which is consistent with that for BI3802 (Figure 3A). Both **D3** and BI3802-induced degradation of BCL6 can be attenuated by the BCL6 inhibitor BI3812, indicating that the binding of **D3** with BCL6 indeed mediate its degradation. Subsequent co-immunoprecipitation study showed that **D3** greatly increased the ubiquitination level of BCL6, further confirmed that **D3** could induce ubiquitin degradation of BCL6 (Figure 3B). These studies suggested that **D3** binds to BCL6 and used a non-Cullin E3 ubiquitin ligase to degrade it by UPS.

**Figure 3.**
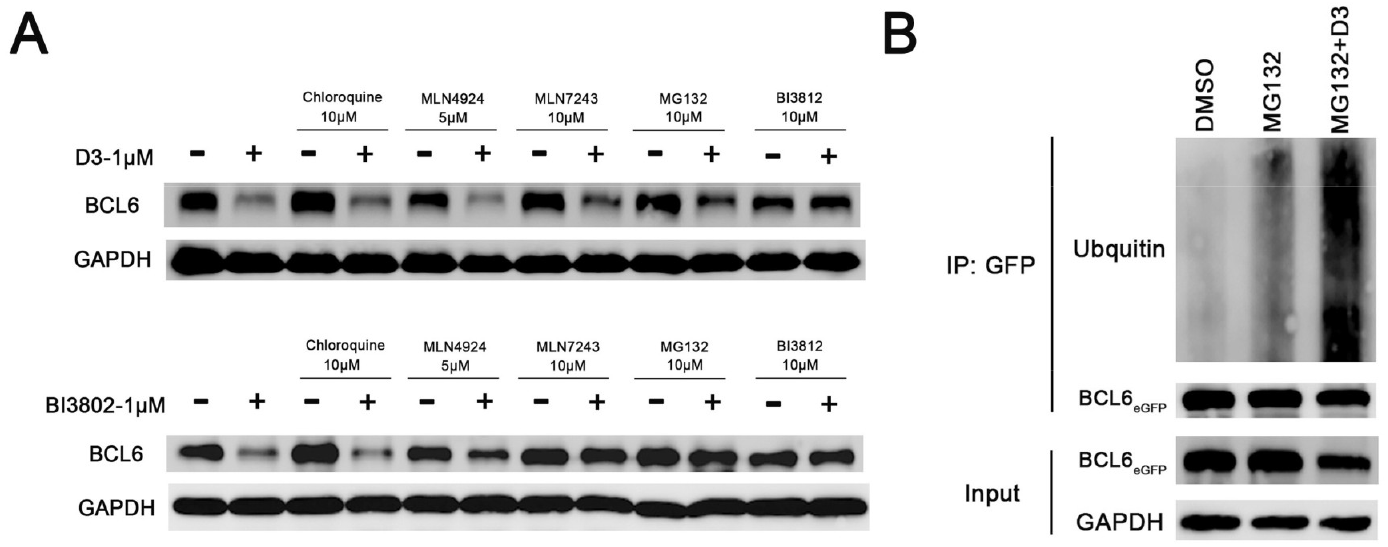
**D3** degrades BCL6 through UPS. A. Immunoblots of BCL6 in Farage cells treated with DMSO, 10 μM MLN7243, 10 μM MG132, 10 μM Chloroquine, 10 μM BI3812, or 5 μM MLN4924 for 15 minutes, then, for indicated samples, 1 μM **D3**/BI3802 was added for next 45 minutes. B. Immunoblots of GFP immunoprecipitation of 10 μM MG132 treatment for 15 min and 1 μM **D3** treatment for next 45min or DMSO/10 μM MG132 treatment for 1 h from HEK293T cells expressing BCL6_eGFP_.

Next, we evaluated the morphological changes of BCL6 protein particles after the addition of **D3** or BI3802 by negative stain electron microscopy. The natural BCL6 protein is present as monodisperse particles as expected, however, both **D3** and BI3802 could obviously induce BCL6 aggregation, although the expected regular helix shape was not observed, which might be attributed to the BCL6 (5-360 aa) in previous study was replaced by the full-length BCL6 in our study^[22]^ (Figure 4A). The hetero-bivalent ligand **D5** couldn’t induce BCL6 aggregation, which explained the reason why it failed to degrade BCL6. Then immunofluorescence study was conducted in Farage cells, endogenous native BCL6 is evenly distributed in the nucleus, the level of BCL6 decreased markedly after **D3** or BI3802 treatments, demonstrating the degradation of BCL6 by both compounds in cells (Figure 4B). The addition of MG132 could substantially reverse the fluorescence decrease caused by **D3** or BI3802, accompanied by the increase of fluorescence foci. Next, in the HEK293T cells stably expressing exogenous BCL6_eGFP_, both **D3** and BI3802 could decrease the level of BCL6_eGFP_, and the number and size of fluorescence foci was also significantly increased after the addition of MG132 (Figure 4C). These results indicated that **D3** could induce the aggregation and degradation of BCL6 like BI3802.

**Figure 4.**
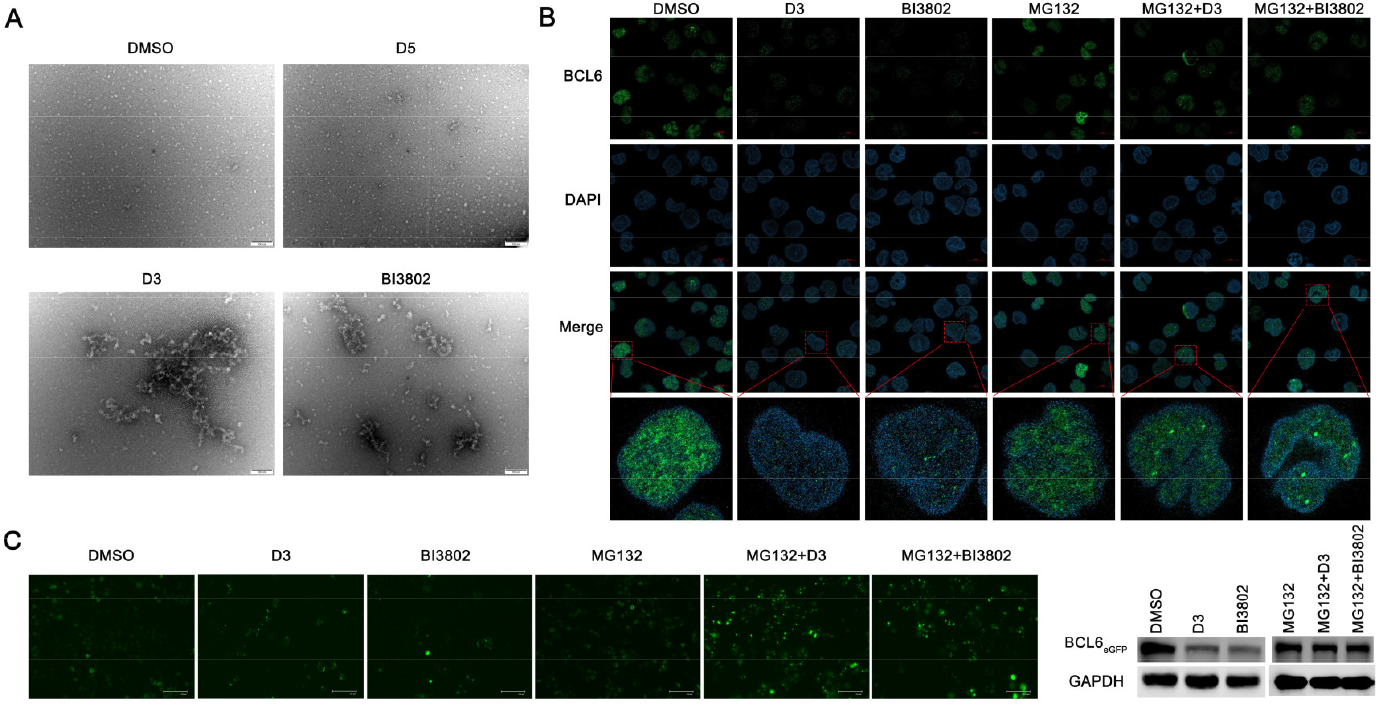
**D3** induced BCL6 polymerized to form cellular foci. A. Negative stain electron microscopy micrographs of BCL6^FL^ protein in DMSO or 12.5 μM **D3**/**D5**/BI3802. Scale bars are 100 nm (n = 3 images). B. Immunofluorescence images of Farage cells treated with DMSO, 1 μM **D3**, 1 μM BI3802, 1 μM MG132, 1 μM MG132 with 1 μM **D3**, or 1 μM MG132 with 1 μM BI3802 for 8 hours. Scale bars are 5 μm (n = 3 images). C. Fluorescence images (left) and immunoblots (right) of HEK293T cells expressing BCL6_eGFP_ after treatment with DMSO, 1 μM **D3**, 1 μM BI3802, 1 μM MG132, 1 μM MG132 with 1 μM **D3**, or 1 μM MG132 with 1 μM BI3802 for 8 hours. Scale bars are 125 μm (n = 3 images).

Since BI3802 degrades BCL6 via its endogenous E3 ligase SIAH1, we speculated that **D3**, which lacks an E3 ligase recruitment fragment may also induce BCL6 degradation through SIAH1. As expected, knocking down of SIAH1 raised the level of BCL6 markedly, indicating that SIAH1 participated in the endogenous degradation of BCL6 as previous study. **D3** also lost the ability to degrade BCL6 after SIAH1 knocked down (Figure 5A). Furthermore, In the presence of **D3** instead of the negative control **D5**, SIAH1 can be more effectively co-immunoprecipitated with BCL6_eGFP_ and colocalized to BCL6 foci in Farage cells, indicating that **D3** could promote the interaction of SIAH1 and BCL6 (Figure 5B, C).

**Figure 5.**
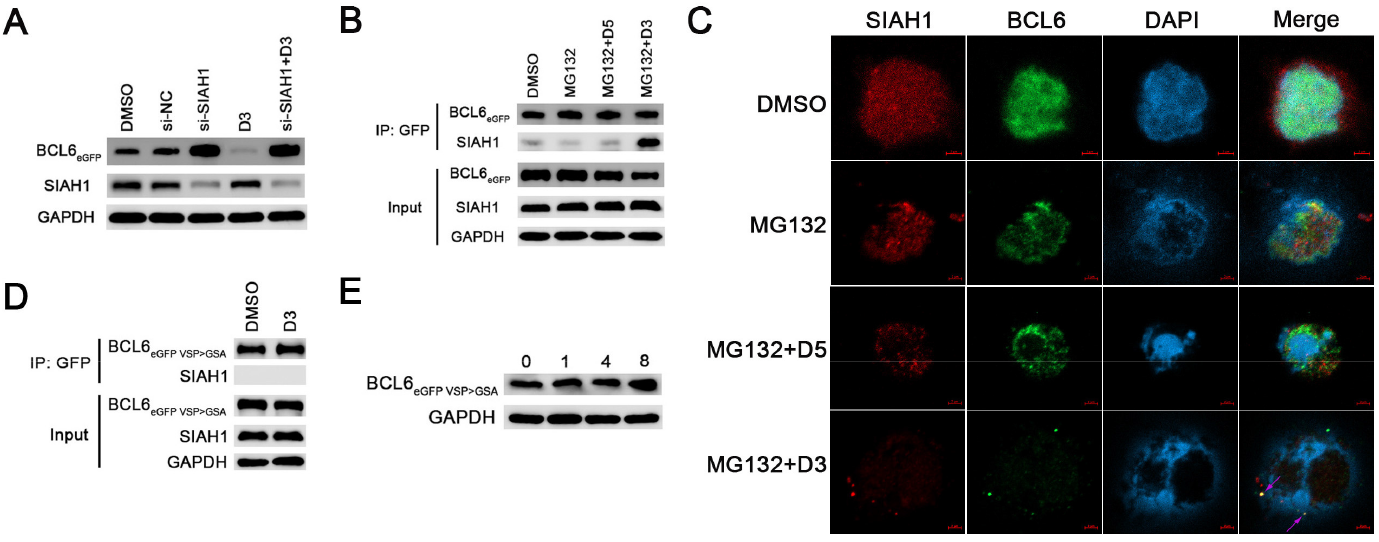
SIAH1 was involved in degradation of polymerized BCL6. A. HEK293T cells expressing BCL6_eGFP_ was transfected by SIAH1 siRNA, siRNA-Negative Control or DMSO for 48 hours, then added 1 μM **D3** to indicated group for 1 hour, detected the levels of BCL6_eGFP_ and SIAH1 by immunoblots. B. Immunoblots of GFP immunoprecipitation of 10 μM MG132 treatment for 15 min and 1 μM **D3** or **D5** treatment for next 45 min or DMSO/10 μM MG132 treatment for 1 h from HEK293T cells expressing BCL6_eGFP_. C. Farage cells were treated with DMSO/1 μM MG132, 1 μM MG132 with 1 μM **D5** or 1 μM MG132 with 1 μM **D3** for 24 h. Cells were imaged by immunofluorescence (n = 3) (the purple arrows refer to the colocalization foci of SIAH1 and BCL6). Scale bars are 2 μm. D. Immunoblots of GFP immunoprecipitation of 1 μM **D3** for 1 h or DMSO from HEK293T cells expressing BCL6_eGFP VSP>GSA_. E. Immunoblots of HEK293T cells expressing BCL6_eGFP VSP>GSA_ after treatment of 1 μM **D3** for 0/1/4/8 hours.

The SIAH1 E3 ligase recognizes a VxP motif on substrate proteins, and this motif is present in the residues 249–251 of BCL6, we mutated the VSP motif of BCL6 to GSA^[20]^. Immunoprecipitation assays showed that SIAH1 was indeed unable to bind to BCL6 after mutation (Figure 5D). As a result, **D3** lost its ability to degrade BCL6 (Figure 5E, Figure S2). Together, these data demonstrated that SIAH1 is an E3 ligase involved in **D3**-induced BCL6 degradation.

We studied the effects of **D3** and BI3802 on BCL6 downstream genes and cell proliferation in Farage cells. **D3**/BI3802 could induced the de-repression of several known BCL6-regulated genes including CCND2, CDKN1A, CD69 and PRDM1 (Figure 6A). Moreover, **D3**/BI3802 leads to a decrease in cell viability (Figure 6B) and proliferative ability (Figure 6C), and can continue to inhibit cell growth even after prolonged exposure (Figure 6D).

**Figure 6.**
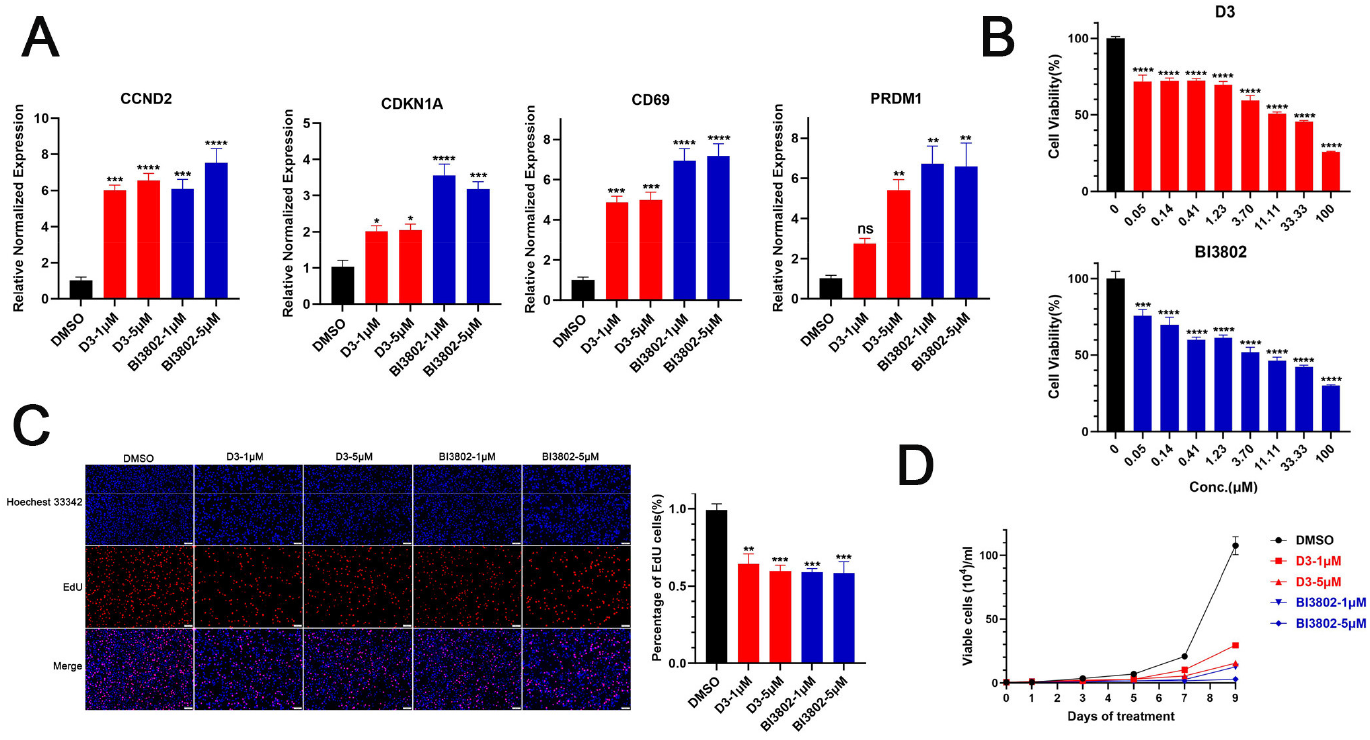
**D3** induced the de-repression of BCL6 target genes and inhibited cell proliferation. A. mRNA levels of CCND2, CDKN1A, CD69 and PRDM1 quantified by qPCR in Farage cells after treatment with 1 or 5 μM **D3**/BI3802 or DMSO for 24 hours. B. Farage cell viability analysis by MTT assays after treatment of **D3**/BI3802 for 3 days.. C. Edu staining was used to detect the farage cell proliferation phenotype with 1 or 5 μM **D3**/BI3802 or DMSO treatment for 48 h, quantification of the EDU analysis was shown on the right of the picture.. D. The quantifications of live Farage cells after 1 or 5 μM **D3**/BI3802 or DMSO treatment, counted the cells every other day and then fitted the proliferation curve. ^ns^p > 0.05. *p < 0.05, **p < 0.01, ***p < 0.001, ****p < 0.0001.

## Conclusions

Herein, we propose and validate protein polymerization induced by rationally designed bivalent ligands as a new means of BCL6 degradation. The BCL6 HBiD **D3** exhibited excellent target selectivity and degradation activity, and can degrade BCL6 rapidly and persistently. This compound can be a good starting point to develop new drug candidate and also be used as a probe to investigate the physiological function of BCL6.

Since protein dimerization is a common phenomenon in cells^[23]^, protein polymerization induced by HBiDs is not limited to BCL6. The abnormal protein polymer induced by HBiDs might be recognized and degraded by protein quality control system including UPS and ALS. Therefore, we speculate that protein degradation caused by polymerization may be a common phenomenon in cells, and HBiDs may be a new type of protein degraders, which is suitable not only for dimerized proteins but also for multi-domain proteins. In fact, some reported HBiD-like molecules, including HCV NS5A inhibitors^[24]^, bivalent inhibitors of BET bromodomains^[25]^ and recent reported intramolecular bivalent glues of BRD4^[26]^, may also exert their biological activity partial by inducing polymerization and degradation of their targets.

Currently reported bivalent ligands for different targets have been proven to be intramolecular ligands, including the recently reported bivalent inhibitors of the BTB E3 ligand KEAP1^[27]^. We believe that the HBiDs reported here are unlikely to be intramolecular bivalent ligands of BCL6 dimers, because their highly rigid structure means that they can only bind to two BTB domains simultaneously when the two BTB domains of a BCL6 dimer are arranged in a head-to-head manner, which is inconsistent with current evidence from structural biology^[28]^. On the other hand, due to differences in modes of action, the morphology of BCL6 polymers induced by HBiDs may be different from that induced by BI3802, which requires subsequent structural biology studies.

Although the development of HBiDs as drugs may also encounter the problem of poor druglike properties faced by other chimeric molecules^[29]^, they can partially avoid the off-target effect of PROTAC as well as its resistance caused by E3 mutations^[30]^. In addition, HBiDs induced protein degradation may be more dependent on the endogenous E3 of POI, it will not be an obstacle to choose a suitable E3 to design HBiDs for POI, such that those proteins which are previously difficult to be targeted by PROTAC could also be degraded by HBiDs.

## Supporting Information

Supplementary figures, supplementary tables, material and methods, NMR and HRMS spectra of compounds.

## Acknowledgements

The authors acknowledge financial support from the Natural Science Foundation of China (No. 22377100), the Natural Science Foundation of Sichuan Province (No. 2022NSFSC1338) and Fundamental Research Funds for the Central Universities (No. 2682023ZTPY011). We would like to thank Analysis and Testing Center of Southwest Jiaotong University for ^1^H-NMR and ^13^C-NMR measurements.

## Reference

[1] J. Salami, C. M. Crews, Science 2017, 355, 1163–1167.

[2] L. Zhao, J. Zhao, K. Zhong, A. Tong, D. Jia, Signal Transduction and Targeted Therapy 2022, 7, 113.

[3] V. Němec, M. P. Schwalm, S. Müller, S. Knapp, Chemical Society Reviews 2022, 51, 7971–7993.

[4] M. Békés, D. R. Langley, C. M. Crews, Nature Reviews Drug Discovery 2022, 21, 181–200.

[5] T. K. Neklesa, H. S. Tae, A. R. Schneekloth, M. J. Stulberg, T. W. Corson, T. B. Sundberg, K. Raina, S. A. Holley, C. M. Crews, Nature Chemical Biology 2011, 7, 538–543.

[6] G. Dong, Y. Ding, S. He, C. Sheng, Journal of Medicinal Chemistry 2021, 64, 10606–10620.

[7] G. Ahn, S. M. Banik, C. L. Miller, N. M. Riley, J. R. Cochran, C. R. Bertozzi, Nature Chemical Biology 2021, 17, 937–946.

[8] S. M. Banik, K. Pedram, S. Wisnovsky, G. Ahn, N. M. Riley, C. R. Bertozzi, Nature 2020, 584, 291–297.

[9] D. Takahashi, H. Arimoto, Autophagy 2020, 16, 765–766.

[10] D. Takahashi, J. Moriyama, T. Nakamura, E. Miki, E. Takahashi, A. Sato, T. Akaike, K. Itto-Nakama, H. Arimoto, Molecular Cell 2019, 76, 797-810.e710.

[11] Z. Li, C. Zhu, Y. Ding, Y. Fei, B. Lu, Autophagy 2019, 16, 185–187.

[12] C. H. Ji, H. Y. Kim, M. J. Lee, A. J. Heo, D. Y. Park, S. Lim, S. Shin, S. Ganipisetti, W. S. Yang, C. A. Jung, K. Y. Kim, E. H. Jeong, S. H. Park, S. Bin Kim, S. J. Lee, J. E. Na, J. I. Kang, H. M. Chi, H. T. Kim, Y. K. Kim, B. Y. Kim, Y. T. Kwon, Nature Communications 2022, 13, 904.

[13] M. He, C. Cao, Z. Ni, Y. Liu, P. Song, S. Hao, Y. He, X. Sun, Y. Rao, Signal Transduction and Targeted Therapy 2022, 7, 181.

[14] M. Schneider, C. J. Radoux, A. Hercules, D. Ochoa, I. Dunham, L.-P. Zalmas, G. Hessler, S. Ruf, V. Shanmugasundaram, M. M. Hann, P. J. Thomas, M. A. Queisser, A. B. Benowitz, K. Brown, A. R. Leach, Nature Reviews Drug Discovery 2021, 20, 789–797.

[15] R. G. Guenette, S. W. Yang, J. Min, B. Pei, P. R. Potts, Chemical Society Reviews 2022, 51, 5740–5756.

[16] L. Zhang, B. Riley-Gillis, P. Vijay, Y. Shen, Molecular Cancer Therapeutics 2019, 18, 1302–1311.

[17] C. Huang, K. Hatzi, A. Melnick, Nature Immunology 2013, 14, 380–388.

[18] C. Duy, C. Hurtz, S. Shojaee, L. Cerchietti, H. Geng, S. Swaminathan, L. Klemm, S.-m. Kweon, R. Nahar, M. Braig, E. Park, Y.-m. Kim, W.-K. Hofmann, S. Herzog, H. Jumaa, H. P. Koeffler, J. J. Yu, N. Heisterkamp, T. G. Graeber, H. Wu, B. H. Ye, A. Melnick, M. Müschen, Nature 2011, 473, 384–388.

[19] H. S. Madapura, N. Nagy, D. Ujvari, T. Kallas, M. C. L. Kröhnke, S. Amu, M. Björkholm, L. Stenke, P. K. Mandal, J. S. McMurray, M. Keszei, L. S. Westerberg, H. Cheng, F. Xue, G. Klein, E. Klein, D. Salamon, Oncogene 2017, 36, 4619–4628.

[20] M. Slabicki, H. Yoon, J. Koeppel, L. Nitsch, S. S. Roy Burman, C. Di Genua, K. A. Donovan, A. S. Sperling, M. Hunkeler, J. M. Tsai, R. Sharma, A. Guirguis, C. Zou, P. Chudasama, J. A. Gasser, P. G. Miller, C. Scholl, S. Frohling, R. P. Nowak, E. S. Fischer, B. L. Ebert, Nature 2020, 588, 164–168.

[21] S. Duan, L. Cermak, J. K. Pagan, M. Rossi, C. Martinengo, P. F. di Celle, B. Chapuy, M. Shipp, R. Chiarle, M. Pagano, Nature 2012, 481, 90–93.

[22] L. Nitsch, P. Jensen, H. Yoon, J. Koeppel, S. S. R. Burman, E. S. Fischer, C. Scholl, S. Fröhling, M. Słabicki, Cell Reports Methods 2022, 2, 100193.

[23] H. Schweke, M. Pacesa, T. Levin, C. A. Goverde, P. Kumar, Y. Duhoo, L. J. Dornfeld, B. Dubreuil, S. Georgeon, S. Ovchinnikov, D. N. Woolfson, B. E. Correia, S. Dey, E. D. Levy, Cell 2024, 187, 999-1010.e1015.

[24] M. Belema, O. D. Lopez, J. A. Bender, J. L. Romine, D. R. St. Laurent, D. R. Langley, J. A. Lemm, D. R. O’Boyle, J.-H. Sun, C. Wang, R. A. Fridell, N. A. Meanwell, Journal of Medicinal Chemistry 2014, 57, 1643–1672.

[25] M. J. Waring, H. Chen, A. A. Rabow, G. Walker, R. Bobby, S. Boiko, R. H. Bradbury, R. Callis, E. Clark, I. Dale, D. L. Daniels, A. Dulak, L. Flavell, G. Holdgate, T. A. Jowitt, A. Kikhney, M. McAlister, J. Méndez, D. Ogg, J. Patel, P. Petteruti, G. R. Robb, M. B. Robers, S. Saif, N. Stratton, D. I. Svergun, W. Wang, D. Whittaker, D. M. Wilson, Y. Yao, Nature Chemical Biology 2016, 12, 1097–1104.

[26] O. Hsia, M. Hinterndorfer, A. D. Cowan, K. Iso, T. Ishida, R. Sundaramoorthy, M. A. Nakasone, H. Imrichova, C. Schätz, A. Rukavina, K. Husnjak, M. Wegner, A. Correa-Sáez, C. Craigon, R. Casement, C. Maniaci, A. Testa, M. Kaulich, I. Dikic, G. E. Winter, A. Ciulli, Nature 2024, 627, 204–211.

[27] M. Lu, J. Ji, Y. Lv, J. Zhao, Y. Liu, Q. Jiao, T. Liu, Y. Mou, Q. You, Z. Jiang, Cell Chemical Biology 2024, 31, 1188-1202.e1110.

[28] N. Kerres, S. Steurer, S. Schlager, G. Bader, H. Berger, M. Caligiuri, C. Dank, J. R. Engen, P. Ettmayer, B. Fischerauer, G. Flotzinger, D. Gerlach, T. Gerstberger, T. Gmaschitz, P. Greb, B. Han, E. Heyes, R. E. Iacob, D. Kessler, H. Kolle, L. Lamarre, D. R. Lancia, S. Lucas, M. Mayer, K. Mayr, N. Mischerikow, K. Muck, C. Peinsipp, O. Petermann, U. Reiser, D. Rudolph, K. Rumpel, C. Salomon, D. Scharn, R. Schnitzer, A. Schrenk, N. Schweifer, D. Thompson, E. Traxler, R. Varecka, T. Voss, A. Weiss-Puxbaum, S. Winkler, X. Zheng, A. Zoephel, N. Kraut, D. McConnell, M. Pearson, M. Koegl, Cell Reports 2017, 20, 2860–2875.

[29] C. Cantrill, P. Chaturvedi, C. Rynn, J. Petrig Schaffland, I. Walter, M. B. Wittwer, Drug Discovery Today 2020, 25, 969–982.

[30] A. Hanzl, R. Casement, H. Imrichova, S. J. Hughes, E. Barone, A. Testa, S. Bauer, J. Wright, M. Brand, A. Ciulli, G. E. Winter, Nature Chemical Biology 2022, 19, 323–333.

