## Supplementary information for "Bivalent ligands as a new universal chemotype of BCL6 degrader by inducing aggregation"

**Luo-Ting Yu**

Department of Biotherapy, Cancer Center and State Key Laboratory of Biotherapy

West China Hospital, Sichuan University

Chengdu, 610041, China

1. Supplementary figures

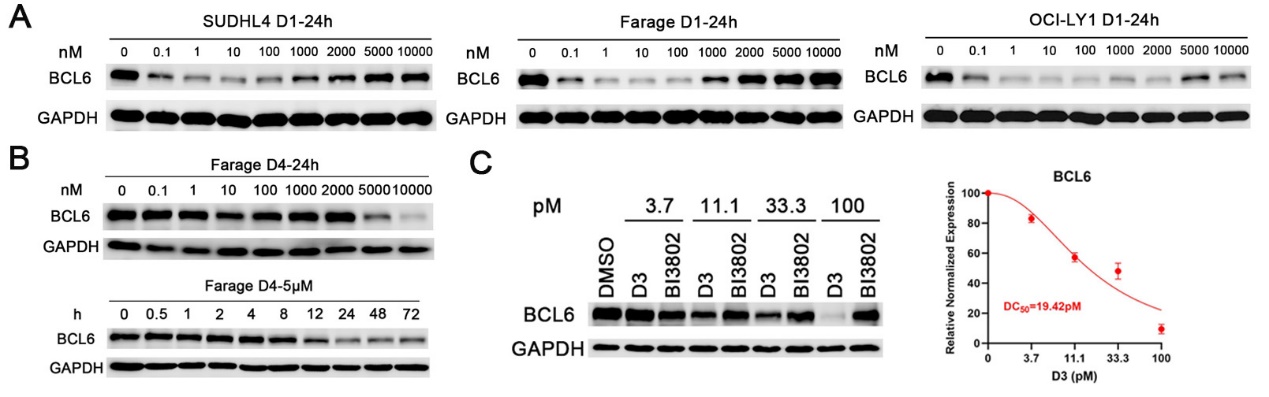

Figure S1. HBiDs degraded BCL6. A. Immunoblotting analysis of protein levels for BCL6 in Farage cells after treatment with D1 for different concentrations. B. Immunoblotting analysis of protein levels for BCL6 in Farage cells after treatment with D4 for different times and concentrations. C. Immunoblotting analysis of protein levels for BCL6 in Farage cells after treatment with different concentrations of D3/BI3802 for 24 h.

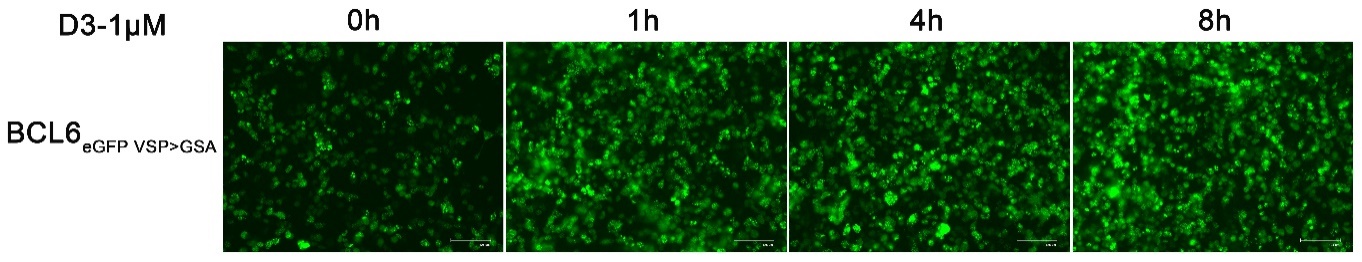

Figure S2. SIAH1 was involved in degradation of polymerized BCL6. Fluorescence images of HEK293T cells expressing BCL6_eGFP VSP>GSA_ after treatment of 1 μM D3 for 0/1/4/8 hours. Scale bars are 125 μm (n = 3 images).

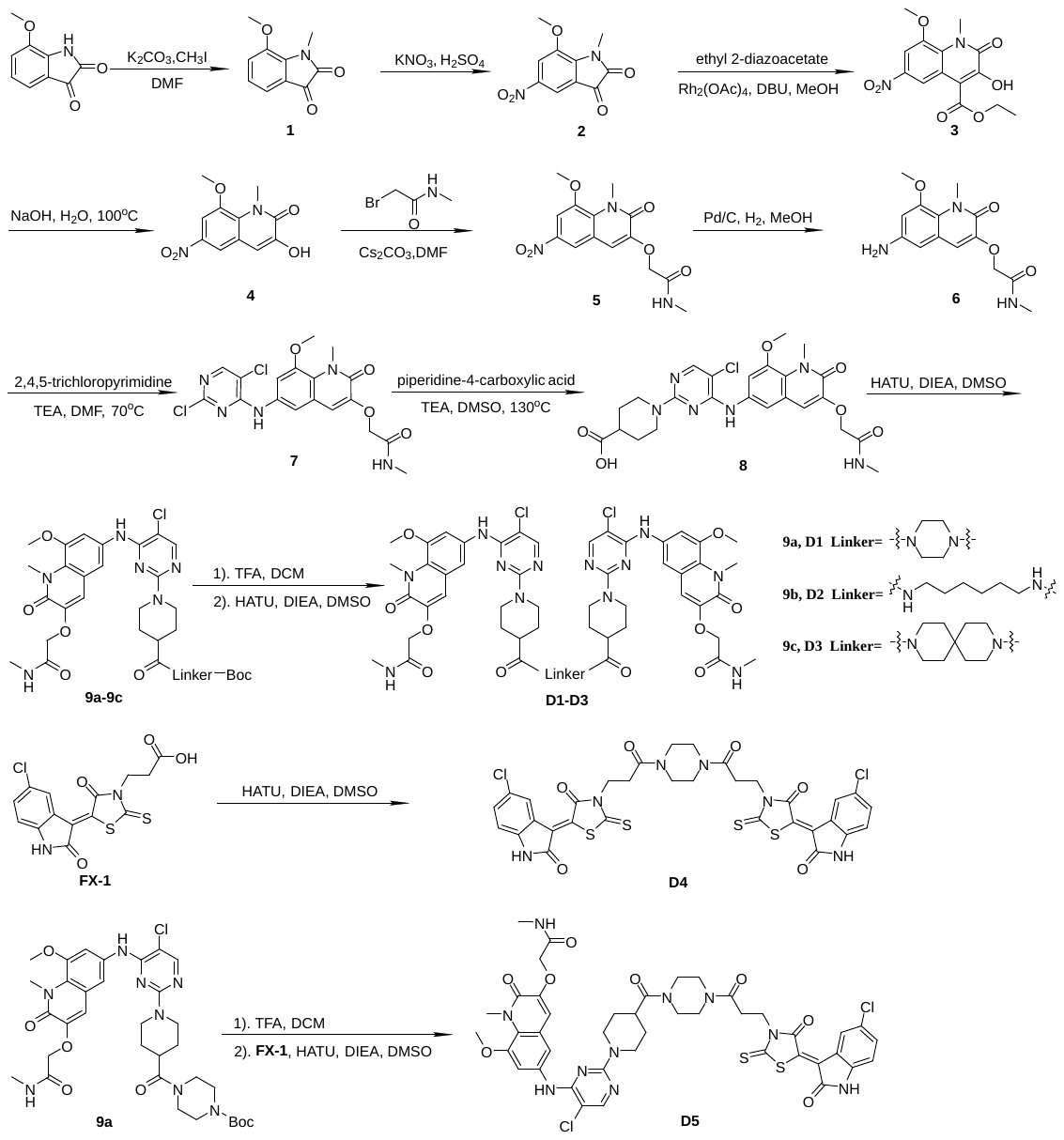

#### Figure S3. Synthesis of bivalent BCL6 ligands D1-D5.

### 2. Supplementary tables

#### Supplementary Tab. S1. The list of antibodies and related information in this article

| **Antibody** | **Company** |
| --- | --- |
| GAPDH | Abway technology |
| BCL6 | CST |
| SIAH1 | Abclonal |
| GFP | Abclonal |
| Ubquitin | Abclonal |

#### Supplementary Tab. S2. The primer sequences used in RT-qPCR

| **Primer** | **Sequences** |
| --- | --- |
| BCL6 | Forward: CTAGGAAAGGCCGGACACC |
|  | Reverse: TGGGAGAGACGTGGGACTAA |
| GAPDH | Forward: ACGGATTTGGTCGTATTGG |
|  | Reverse: TCCCGTTCTCAGCCTTG |
| CCND2 | Forward: GCAGAAGGACATCCAACCCT |
|  | Reverse: GTCGGTGTAAATGCACAGCTT |
| CDKN1A | Forward: GCGACTGTGATGCGCTAATG |
|  | Reverse: GAAGGTAGAGCTTGGGCAGG |
| CD69 | Forward: AAGTTCCTGTCCTGTGTGCT |
|  | Reverse: AAACATGGCTGTCTGATGGC |
| PRDM1 | Forward: TTGTGTGGTATTGTCGGGACTTT |
|  | Reverse: TTTCTCAGTGCTCGGTTGCTTTA |

### 3. Material and Methods

#### 3.1 Compounds synthesis

##### **3.1.1 Materials and general procedures**

For chemical synthesis, all materials were obtained from commercial suppliers and used without further puriﬁcation. The ^1^H and ^13^C NMR spectra were collected on a Bruker Avance 400 or Bruker Avance 600 spectrometer at 25 ℃ using CDCl_3_ or DMSO-*d*_6_ as the solvent. Chemical shifts (*δ*) are reported in ppm relative to TMS (internal standard), coupling constants (*J*) are reported in hertz, and peak multiplicity are reported as s (singlet), d (doublet), t (triplet), q (quartet), m (multiplet), or br s (broad singlet). Mass spectra (MS) were measured on a Micromass Q-TOF Premier mass spectrometer with electron spray ionization (ESI). High resolution mass spectra analysis is performed on a Shimadzu LCMS IT-TOF Chromatography/Mass Spectrometer with electron spray ionization (ESI). Thin layer chromatography (TLC) was performed on 0.20 mm silica gel F-254 plates (Qingdao Haiyang Chemical, China). Preparative thin layer chromatography was performed on 1.00 mm silica gel F-254 plates (Qingdao Haiyang Chemical, China). Visualization of TLC was accomplished with UV light and/or I_2_ in silica gel. Column chromatography was performed using silica gel of 300-400 mesh (Qingdao Haiyang Chemical, China).

##### 3.1.2 Synthesis of bivalent BCL6 ligands D1-D5

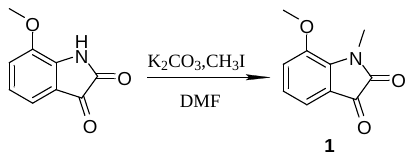

**Synthesis of 7-methoxy-1-methylindoline-2,3-dione (1).**

To a solution of 7-methoxyindoline-2,3-dione (3.54 g, 20 mmol, 1 eq) in DMF (8 mL) was added K_2_CO_3_ (4.14 g, 30 mmol, 1.5 eq). The mixture was stirred at room temperature for 30 min. Then CH_3_I (2 mL, 26 mmol, 1.3 eq) was added and the reaction mixture was stirred at room temperature for 2 h. The resulting solution was poured into water (50 mL) and acidized with 4 M HCl after the reaction completed, then it was ﬁltered, and the residue was washed with water and dried under vacuum to give 7-methoxy-1-methylindoline-2,3-dione (**1**, 3.134g, 16.41 mmol) as a solid, yield 82 %. ^1^H NMR (400 MHz, Chloroform-*d*) *δ* 7.22 (dd, *J* = 7.4, 1.1 Hz, 1H), 7.16 (dd, *J* = 8.4, 1.1 Hz, 1H), 7.05 (dd, *J* = 8.3, 7.3 Hz, 1H), 3.90 (s, 3H), 3.50 (s, 3H).

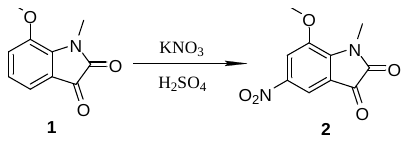

**Synthesis of 7-methoxy-1-methyl-5-nitroindoline-2,3-dione (2)**.

To a solution of **1** (3.441 g, 18 mmol, 1 eq) in concentrated sulfuric acid (20 mL) was added KNO_3_ (1.74 g, 18 mmol, 1 eq)，and the mixture was stirred at 0 °C for 30 min. Then the reaction mixture was allowed to stir at room temperature for 18 h. Then another part of KNO_3_ (435 mg, 4.5 mmol, 0.25 eq) was added and the reaction mixture was stirred at room temperature for another 1 h until the starting material was completely converted as determined by TLC. The resulting mixture was poured into ice water and extracted with ethyl acetate. The organic phase was washed with brine and dried over anhydrous Mg_2_SO_4_, followed by removing of the solvent to afford the crude product, which was purified by silica gel chromatography to give 7-methoxy-1-methyl-5-nitroindoline-2,3-dione (**2**, 3.099 g, 13.12 mmol) as a red solid, yield 73 %. ^1^H NMR (400 MHz, DMSO-*d*_6_) *δ* 8.10 (d, *J* = 2.2 Hz, 1H), 7.90 (d, *J* = 2.2 Hz, 1H), 4.01 (s, 3H), 3.40 (s, 3H).

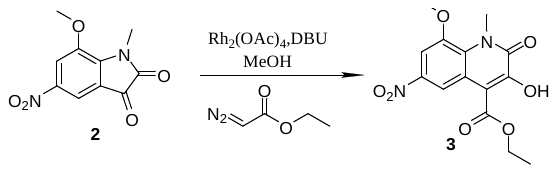

**Synthesis of 3-hydroxy-8-methoxy-1-methyl-6-nitro-2-oxo-1,2-dihydroquinoline-4-carboxylate (3)**.

To a solution of **2** (1.425 g, 6 mmol, 1 eq) in MeOH (60 mL) was added DBU (180 μL, 1.2 mmol, 0.2 eq) and ethyl 2-diazoacetate (1.5 mL, 12 mmol, 2 eq), and the mixture was stirred at room temperature for 1 h. Dirhodiumtetraacetate (51 mg, 0.12 mmol, 0.02 eq) was added and the reaction mixture was stirred at room temperature for another 12 h. Then the resulting mixture was ﬁltered, and the residue was washed with MeOH and dried under vacuum to give ethyl 3-hydroxy-8-methoxy-1-methyl-6-nitro-2-oxo-1,2-dihydroquinoline-4-carboxylate (**3**, 1.557 g, 4.83 mmol) as a yellow solid, yield 80 %. ^1^H NMR (400 MHz, DMSO-*d*_6_) *δ* 8.01 (s, 1H), 7.55 (s, 1H), 4.32 (q, *J* = 7.1 Hz, 2H), 3.97 (s, 2H), 3.85 (s, 3H), 1.28 (t, *J* = 7.1 Hz, 3H).

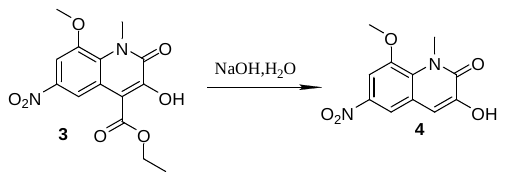

**Synthesis of 3-hydroxy-8-methoxy-1-methyl-6-nitroquinolin-2(1H)-one (4)**.

To a solution of **3** (645 mg, 2 mmol, 1 eq) in H_2_O (20 mL) was added NaOH (1.600 g, 40 mmol, 20 eq) , and the mixture was stirred at 100 °C for 16 h until the starting material was completely converted as determined by TLC. The resulting mixture was cooled to room temperature and it was acidized with 2 M HCl to pH 2−3. After filtration, the residue was washed with water and dried under vacuum to give 3-hydroxy-8-methoxy-1-methyl-6-nitroquinolin-2(1H)-one (**4**, 431 mg, 1.72 mmol) as a yellow solid, yield 86 %. ^1^H NMR (400 MHz, DMSO-*d*_6_) *δ* 10.03 (s, 1H), 8.20 (d, *J* = 2.4 Hz, 1H), 7.73 (d, *J* = 2.4 Hz, 1H), 7.31 (s, 1H), 4.00 (s, 3H), 3.92 (s, 3H).

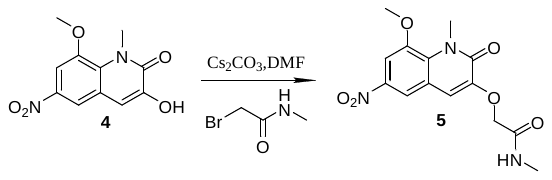

**Synthesis of 2-((8-methoxy-1-methyl-6-nitro-2-oxo-1,2-dihydroquinolin-3-yl)oxy)-N-methylacetamide (5).**

To a solution of **4** (250 mg, 1 mmol, 1 eq) in DMF was added 2-bromo-N-methylacetamide (180 mg,1.2 mmol, 1.2 eq) and Cs_2_CO_3_ (650 mg, 2 mmol, 2 eq), and the mixture was stirred at room temperature for 1 h. The reaction solution was poured into water after the starting material was completely converted as determined by TLC. Then the resulting mixture was ﬁltered, and the residue was washed with water and dried under vacuum to give 2-((8-methoxy-1-methyl-6-nitro-2-oxo-1,2-dihydroquinolin-3-yl)oxy)-N-methylacetamide (**5**, 229 mg, 0.71 mmol) as a yellow solid, yield 71 %. ^1^H NMR (400 MHz, DMSO-*d*_6_) *δ* 8.24 (d, *J* = 2.5 Hz, 1H), 7.91 (s, 1H), 7.79 (d, *J* = 2.5 Hz, 1H), 7.46 (s, 1H), 4.58 (s, 2H), 4.01 (s, 3H), 3.89 (s, 3H), 2.66 (d, *J* = 4.5 Hz, 3H).

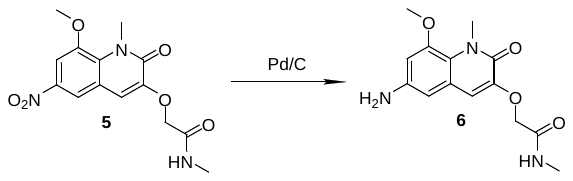

**Synthesis of 2-((6-amino-8-methoxy-1-methyl-2-oxo-1,2-dihydroquinolin-3-yl)oxy)-N-methylacetamide (6)**

To a solution of **5** (321 mg, 1 mmol, 1 eq) in MeOH (20 mL) was added 10% Pd/C (106 mg, 0.1 mmol, 0.1 eq) , and the reaction mixture was stirred at room temperature in a hydrogen atmosphere for 2 h. When the reaction was deemed complete as determined by TLC, the resulting mixture was ﬁltered through diatomaceous earth. The filtrate was concentrated under vacuum to get the crude product, which was purified by silica gel chromatography to give 2-((6-amino-8-methoxy-1-methyl-2-oxo-1,2-dihydroquinolin-3-yl)oxy)-N-methylacetamide (**6**, 192 mg, 0.66 mmol) as a yellow solid , yield 66 %. ^1^H NMR (400 MHz, DMSO-*d*_6_) *δ* 7.92 (d, *J* = 5.3 Hz, 1H), 6.95 (s, 1H), 6.47 (d, *J* = 2.3 Hz, 1H), 6.27 (d, *J* = 2.3 Hz, 1H), 5.08 (s, 2H), 4.49 (s, 2H), 3.79 (s, 3H), 3.77 (s, 3H), 2.66 (d, *J* = 4.7 Hz, 3H).

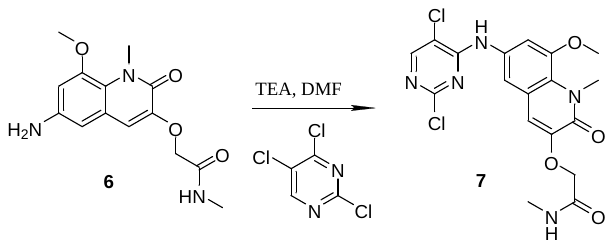

**Synthesis of 2-((6-((2,5-dichloropyrimidin-4-yl)amino)-8-methoxy-1-methyl-2-oxo-1,2-dihydroquinolin-3-yl)oxy)-N-methylacetamide (7)**

To a solution of **6** (145 mg, 0.5 mmol, 1 eq) in DMF (5 mL) was added TEA (248 μL, 1.5 mmol, 3 eq) and 2,4,5-trichloropyrimidine (92 mg, 0.5 mmol, 1 eq) , and the mixture was stirred at 70 °C for 16 h. When the reaction was deemed complete as determined by TLC, the reaction mixture was cooled to room temperature. Then the resulting solution was ﬁltered, and the residue was washed with water and dried under vacuum to give 2-((6-((2,5-dichloropyrimidin-4-yl)amino)-8-methoxy-1-methyl-2-oxo-1,2-dihydroquinolin-3-yl)oxy)-N-methylacetamide (**7**, 182 mg, 0.42 mmol) as a gray solid, yield 83 %. ^1^H NMR (400 MHz, Chloroform-*d*) *δ* 8.22 (s, 1H), 7.65 (s, 1H), 7.35 (s, 2H), 7.00 (s, 1H), 4.50 (s, 2H), 4.05 (d, *J* = 5.0 Hz, 3H), 3.98 (s, 3H), 2.91 (s, 3H).

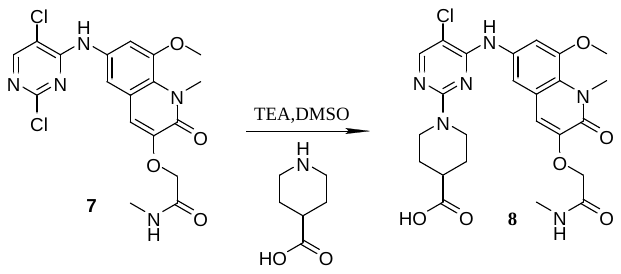

**Synthesis of 1-(5-chloro-4-((8-methoxy-1-methyl-3-(2-(methylamino)-2-oxoethoxy)-2-oxo-1,2-dihydroquinolin-6-yl)amino)pyrimidin-2-yl)piperidine-4-carboxylic acid (8)**

To a solution of **7** (132 mg, 0.3 mmol, 1 eq) in DMSO (3 mL) was added TEA (166 μL, 1.2 mmol, 4 eq) and piperidine-4-carboxylic acid (46 mg, 0.36 mmol, 1.2 eq) , and the mixture was stirred at 130 °C for 12 h. When the reaction was deemed complete as determined by TLC, the reaction solution was cooled to room temperature and it was acidized with 4 M HCl. Then the resulting mixture was ﬁltered, and the residue was washed with water and dried under vacuum to give 1-(5-chloro-4-((8-methoxy-1-methyl-3-(2-(methylamino)-2-oxoethoxy)-2-oxo-1,2-dihydroquinolin-6-yl)amino)pyrimidin-2-yl)piperidine-4-carboxylic acid (**8**, 123 mg, 0.23 mmol) as a gray solid, yield 77 %. ^1^H NMR (400 MHz, DMSO-*d*_6_) *δ* 8.77 (s, 1H), 8.06 (s, 1H), 7.98 (d, *J* = 4.8 Hz, 1H), 7.54 (s, 2H), 7.00 (s, 1H), 4.55 (s, 2H), 4.45 – 4.35 (m, 2H), 3.88 (s, 3H), 3.86 (s, 3H), 3.07 – 2.97 (m, 2H), 2.66 (d, *J* = 4.6 Hz, 3H), 2.53 (d, *J* = 11.9 Hz, 1H), 1.85 (dd, *J* = 13.5, 3.6 Hz, 2H), 1.54 – 1.44 (m, 2H). ^13^C NMR (100 MHz, DMSO) *δ* 176.27, 167.94, 159.52, 158.22, 155.68, 155.12, 148.07, 147.05, 134.80, 122.84, 121.88, 114.12, 112.61, 107.41, 102.49, 68.35, 57.06, 49.06, 46.01, 43.68, 40.88, 35.49, 27.95, 25.85.

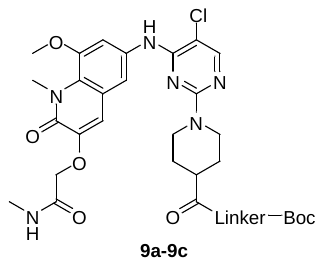

**General procedure for the synthesis of 9a-9c.** To a solution of **8** (159 mg, 0.3 mmol, 1 eq) in DMSO (3 mL) was added N-Boc diamine (0.36 mmol, 1.2 eq), HATU (228 mg, 0.6 mmol, 2 eq) and DIEA (208 μL, 1.20 mmol, 4 eq), and the mixture was stirred at room temperature for 6 h. When the reaction was deemed complete as determined by TLC, the resulting mixture was poured into water and it was ﬁltered. The residue was washed with water and dried under vacuum to afford the crude products **9a-9c**, which were used in the next step without further purification.

**General procedure for the synthesis of D1-D3**

To a solution of compound **9a-9c** (0.3 mmol, 1 eq) in DCM (10 mL) was added TFA (5 mL), and the reaction mixture was stirred at room temperature for 2 h. When the reaction was deemed complete as determined by TLC, the resulting solution was dried under vacuum to give the corresponding amine trifluoroacetate salts as a yellowish semisolid.

Then these amine trifluoroacetate salts (0.1 mmol, 1 eq) were dissolved in DMSO (3 mL) and compound **8** was added, after by the addition of HATU (76 mg, 0.2 mmol, 2 eq) and DIEA (86 μL, 0. 5 mmol, 5 eq). The mixture was stirred at room temperature for 6 h. When the reaction was deemed complete as determined by TLC, the resulting solution was ﬁltered, and the residue was washed and dried under vacuum to afford the crude product, which was purified by silica gel chromatography to give **D1-D3** as a solid, with the total yield in the three steps of 20%-30%.

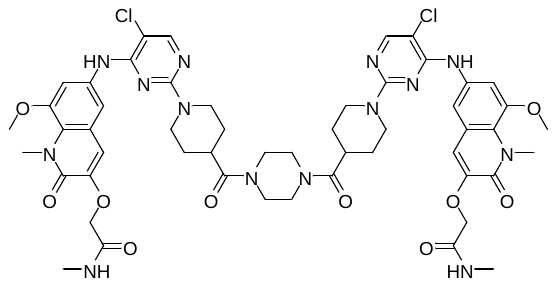

**2,2'-((((((piperazine-1,4-dicarbonyl)bis(piperidine-4,1-diyl))bis(5-chloropyrimidine-2,4-diyl))bis(azanediyl))bis(8-methoxy-1-methyl-2-oxo-1,2-dihydroquinoline-6,3-diyl))bis(oxy))bis(N-methylacetamide) (D1)**

^1^H NMR (600 MHz, DMSO-*d*_6_, T=80 ℃) *δ* 8.62 (s, 2H), 8.06 (s, 2H), 7.82 (s, 2H), 7.58 (s, 2H), 7.55 (s, 2H), 7.04 (s, 2H), 4.53 (s, 8H), 3.89 (d, *J* = 12.9 Hz, 12H), 3.55 (s, 8H), 3.06 – 2.90 (m, 6H), 2.68 (s, 6H), 1.71 (d, *J* = 12.8 Hz, 4H), 1.56 (t, *J* = 12.1 Hz, 4H).

^13^C NMR (150 MHz, DMSO-*d*_6_) δ 173.24, 168.00, 159.68, 158.40, 155.82, 155.13, 148.20, 147.15, 134.90, 123.20, 121.96, 114.87, 112.86, 107.94, 102.56, 68.79, 57.33, 43.86, 40.80, 37.77, 35.35, 28.33, 25.80.

HRMS (ESI)^-^: calcd for C_52_H_59_Cl_2_N_14_O_10_ [M - H]^-^ : 1109.3921, found: 1109.3816.

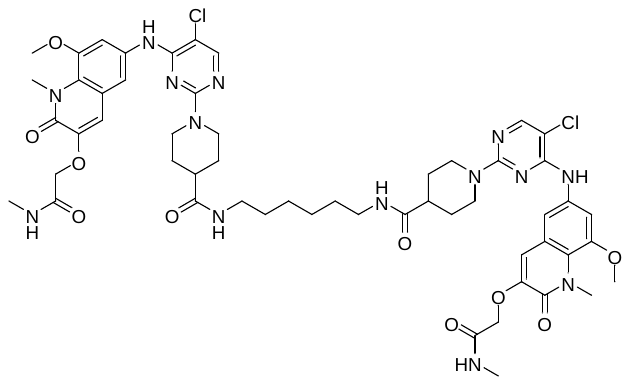

**N,N'-(hexane-1,6-diyl)bis(1-(5-chloro-4-((8-methoxy-1-methyl-3-(2-(methylamino)-2-oxoethoxy)-2-oxo-1,2-dihydroquinolin-6-yl)amino)pyrimidin-2-yl)piperidine-4-carboxamide) (D2)**

^1^H NMR (600 MHz, DMSO-*d*_6_, T=80 ℃) δ 9.37 (s, 0.3H), 8.50 (s, 1.5H), 8.36 (s, 0.2H), 8.03 (s, 1H), 7.79 (s, 2H), 7.59 – 7.52 (m, 3.2H), 7.52 – 7.45 (m, 2.5H), 7.18 (s, 0.5H), 7.05 (s, 1.7H), 4.60 – 4.45 (m, 8H), 3.97 – 3.84 (m, 12H), 3.22 – 3.07 (m, 4H), 3.00 – 2.89 (m, 4H), 2.70 (dd, *J* = 7.8, 4.7 Hz, 6H), 2.43 – 2.33 (m, 2H), 1.79 – 1.68 (m, 4H), 1.59 – 1.48 (m, 4H), 1.43 – 1.34 (m, 4H), 1.30 – 1.24 (m, 4H).

^13^C NMR (150 MHz, DMSO-*d*_6_) δ 179.09, 172.70, 164.19, 163.02, 160.44, 159.74, 152.84, 151.68, 139.59, 127.56, 126.65, 118.88, 107.20, 73.07, 61.81, 48.74, 47.33, 43.44, 40.35, 40.27, 34.22, 33.24, 31.15, 30.59.

HRMS (ESI)^+^: calcd for C_54_H_66_Cl_2_N_14_NaO_10_ [M + Na]^+^ : 1163.4356, found: 1163.4335.

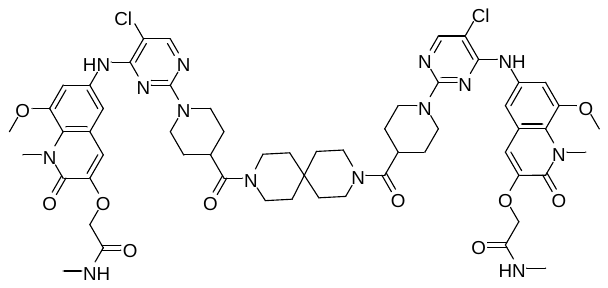

**2,2'-((((((3,9-diazaspiro[5.5]undecane-3,9-dicarbonyl)bis(piperidine-4,1-diyl))bis(5-chloropyrimidine-2,4-diyl))bis(azanediyl))bis(8-methoxy-1-methyl-2-oxo-1,2-dihydroquinoline-6,3-diyl))bis(oxy))bis(N-methylacetamide) (D3)**

^1^H NMR (600 MHz, DMSO-*d*_6_) δ 8.77 (s, 2H), 8.05 (s, 2H), 8.02 (t, *J* = 4.8 Hz, 2H), 7.56 (d, *J* = 2.2 Hz, 2H), 7.53 (d, *J* = 2.3 Hz, 2H), 7.00 (s, 2H), 4.53 (d, *J* = 17.1 Hz, 8H), 3.87 (d, *J* = 9.4 Hz, 12H), 3.48 (d, *J* = 31.6 Hz, 8H), 2.94 (dt, *J* = 19.3, 11.4 Hz, 6H), 2.66 (d, *J* = 4.6 Hz, 6H), 1.65 (d, *J* = 12.4 Hz, 4H), 1.49 (d, *J* = 16.4 Hz, 8H), 1.38 (s, 4H).

^13^C NMR (150 MHz, DMSO) δ 172.65, 167.93, 159.54, 158.25, 155.68, 155.13, 148.10, 146.94, 134.86, 122.78, 121.88, 114.13, 112.62, 107.39, 102.42, 68.35, 57.07, 43.85, 41.05, 37.76, 37.33, 36.33, 35.52, 35.00, 30.78, 28.43, 25.85.

HRMS (ESI)^+^: calcd for C_57_H_68_Cl_2_N_14_NaO_10_ [M + Na]^+^ : 1201.4512, found: 1201.4508

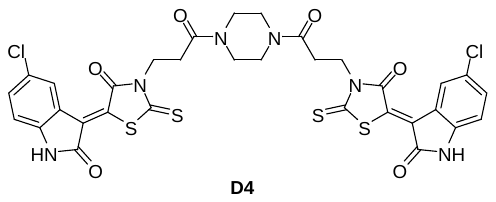

**Synthesis of (Z)-3,3'-(piperazine-1,4-diylbis(3-oxopropane-3,1-diyl))bis(5-((Z)-5-chloro-2-oxoindolin-3-ylidene)-2-thioxothiazolidin-4-one) (D4)**

To a solution of **FX-1** (184 mg, 0.5 mmol, 2 eq) in DMF (5 mL) was added piperazine (22 mg, 0.25 mmol, 1 eq), HATU (285 mg, 3 eq) and DIEA (175 μL, 4 eq), respectively. The mixture was stirred at room temperature for 6 h until the reaction was deemed complete as determined by TLC. The resulting mixture was poured into water and ﬁltered, and the residue was washed with water and dried under vacuum to afford the crude product, which was purified by silica gel chromatography to give **D4** (63 mg, 0.08 mmol) as a red solid, yield 32 %.

^1^H NMR (400 MHz, DMSO-*d*_6_) *δ* 11.42 (s, 2H), 8.84 (d, *J* = 2.2 Hz, 2H), 7.50 (dd, *J* = 8.4, 2.2 Hz, 2H), 7.00 (d, *J* = 8.4 Hz, 2H), 4.32 – 4.24 (m, 4H), 3.48 (s, 8H), 2.77 (t, *J* = 7.9 Hz, 4H).

^13^C NMR (100 MHz, DMSO) *δ* 197.55 (2C), 168.94, 168.60, 168.22, 167.20 , 143.82, 133.31, 132.81, 127.45, 126.49, 124.36, 121.54, 112.71, 45.93, 45.64, 45.26, 44.94, 41.21, 40.81, 21.67.

HRMS (ESI)^-^: calcd for C_32_H_23_Cl_2_N_6_O_6_S_4_ [M + H]^+^ : 784.9944, found: 784.9828

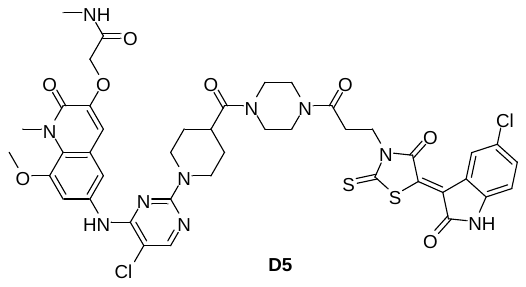

**Synthesis of (Z)-2-((6-((5-chloro-2-(4-(4-(3-(5-(5-chloro-2-oxoindolin-3-ylidene)-4-oxo-2-thioxothiazolidin-3-yl)propanoyl)piperazine-1-carbonyl)piperidin-1-yl)pyrimidin-4-yl)amino)-8-methoxy-1-methyl-2-oxo-1,2-dihydroquinolin-3-yl)oxy)-N-methylacetamide (D5)**

To a solution of compound **9a** (70 mg, 0.1 mmol, 1 eq) in DCM (5 mL) was added TFA (2.5 mL), and the reaction mixture was stirred at room temperature for 2 h. When the reaction was deemed complete as determined by TLC, the resulting solution was dried under vacuum to give 2-((6-((5-chloro-2-(4-(piperazine-1-carbonyl)piperidin-1-yl)pyrimidin-4-yl)amino)-8-methoxy-1-methyl-2-oxo-1,2-dihydroquinolin-3-yl)oxy)-N-methylacetamide trifluoroacetate salt as a yellowish semisolid.

Subsequently, the above-mentioned trifluoroacetate salt were dissolved in DMSO (3 mL) and **FX-1** was added, after by the addition of HATU (76 mg, 0.2 mmol, 2 eq) and DIEA (86 μL, 0.5 mmol, 5 eq). The mixture was stirred at room temperature for 6 h. When the reaction was deemed complete as determined by TLC, the resulting solution was ﬁltered, and the residue was washed and dried under vacuum to afford the crude product, which was purified by silica gel chromatography to give **D5** (22 mg, 0.023 mmol) as a solid, yield 23 %.

^1^H NMR (600 MHz, DMSO-*d*_6_) δ 9.07 (d, *J* = 25.7 Hz, 1H), 8.79 (s, 1H), 8.06 (s, 1H), 8.03 (d, *J* = 4.9 Hz, 1H), 7.58 (s, 1H), 7.55 – 7.51 (m, 1H), 7.50 – 7.40 (m, 1H), 7.08 – 6.95 (m, 2H), 4.53 (d, *J* = 12.0 Hz, 4H), 4.12 (s br, 2H), 3.87 (d, *J* = 9.8 Hz, 6H), 3.76 (s, 2H), 3.63 – 3.43 (m, 6H), 2.97 (t, *J* = 7.9 Hz, 3H), 2.70 – 2.57 (m, 5H), 1.67 (d, *J* = 12.3 Hz, 2H), 1.50 (s br, 2H).

^13^C NMR (150 MHz, DMSO) δ 173.18, 169.17, 167.93, 166.27, 165.80, 165.18, 159.53, 158.26, 155.71, 155.14, 148.12, 146.94, 134.87, 130.19, 128.90, 128.62, 127.67, 122.80, 121.89, 117.37, 114.13, 112.65, 107.47, 102.47, 68.34, 57.10, 55.35, 45.08, 43.81, 40.61, 37.71, 35.51, 28.30, 25.84, 21.54.

HRMS (ESI)^-^: calcd for C_42_H_41_Cl_2_N_10_O_8_S_2_ [M - H]^-^ : 947.1933, found: 947.1848

#### 3.2 Cell culture

All the cell lines used in this study were purchased from the American Type Culture Collection (ATCC, USA) or the Cell Bank of the Chinese Academy of Science (Shanghai, China). DLBCL cells (Farage, SUDHL4, OCI-LY1) were cultured in Rosewell Park Memorial Institute (RPMI) 1640 media and HEK293T cells were cultured in Dulbecco’s modified Eagle’s medium (DMEM), with 1% penicillin-streptomycin and 10% fetal bovine serum (FBS) at 37 ℃ and 5% CO_2_.

#### 3.3 Western blot analysis

After treatment of indicated compounds and times, cells were harvested and lysed in RIPA buffer (Beyotime, China) containing cocktail (1:100, MCE, USA) on ice for 30 min. Then the cell lysates were centrifuged at 12,000 rpm at 4 ℃ for 15 min, the supernatant was harvested and the protein concentration was determined by the BCA (Beyotime, China) kit. Protein samples were diluted by RIPA buffer to the same concentration and denatured by 100 ℃ water bath for 10 min after adding loading buffer. The samples were separated on SDS-PAGE gel and transferred onto polyvinylidene fluoride (PVDF) filter membranes (BIO-RAD, USA). Then the membranes were incubated with appropriate primary antibody and corresponding secondary antibody. The immunoreactive protein bands were detected using an enhanced chemiluminescence kit (Oriscience, China). The antibody used in this study is listed in the supplementary table S1.

#### 3.4 Proteomics

The quantitative proteomics analysis was commissioned to be completed by PTM BIO, performed by following the manufacturer’s instruction.

#### 3.5 Real time qPCR assay

Cells were treated with different compounds for indicated time, then total RNA was extracted with trizol. RNA was reverse-transcribed by Hiscript® III RT SuperMix (Vazyme, China). RT-qPCR was carried out using the ChamQ Universal SYBR qPCR Master Mix (Vazyme, China) on the CFX96 RT-qPCR system in accordance with the manufacturer’s instruction. The reaction procedure was as follows: 95 ℃ for 30 s followed by 40 cycles of amplification for 6 s at 95 ℃, 20 s at 60 ℃. The related primer sequences is listed in supplementary table S2.

#### 3.6 Lentivirus Production

In a 6-well plate format, 5 × 10^5^ HEK293T cells were seeded per well in 2 mL medium. The next day, 21.7 ul lipo 3000 transfection reagent (Invitrogen, USA) was added to 500 ul OPTI-MEM (Invitrogen, USA) (mix A). According to the ratio of 4:3:1, BCL6_eGFP_ plasmid (LabLead, China), psPAX2, and pMD.2G were added to 500 ul OPTI-MEM (Invitrogen, USA) (mix B). Adding mix B to mix A and incubated for 15 min at RT then added it to HEK293T cells in a dropwise manner. The supernatant was collected after 60 h, filtered with a 0.45 μm filter, and incubated with 1× PEG-8000 at 4 °C over night. Then centrifuged at 1600 g for 45 min. The precipitation was resuscitated with 5 ml of DMEM medium.

#### 3.7 Lentivirus Infection

2.5 × 10^5^ HEK293T cells were seeded per well in to a 6-well plate format, the next day, changed the medium to 0.5 ml virus solution above and 1.75 ml DMEM medium with 0.1% polybrene. After 48 h infection, changed the medium with 2 ml fresh medium containing 3 ug/ml puromycin and replaced it every 3 days. The selection lasted 10 days.

#### 3.8 Co-immunoprecipitation

HEK293T cells expressing BCL6_eGFP_ were plated into 10 cm dishes, cultured for one day, then treated with indicated compounds and times. The cells were harvested and lysed in NP40 lysis buffer (Beyotime, China) infused with protease inhibitor (MCE, USA) for 15 min on ice. Lysates were cleared by centrifugation (12,000 rpm, 15 min, 4 °C). The lysate was divided into two parts, input and IP. 50 μL of pre-cleaned Protein A/G Magnetic Beads incubated with GFP antibodies for 2 h at 4 °C, was added to the IP lysates. The beads-lysate mix was incubated at 4 °C for 5 h. Proteins were eluted in sample buffer at 95 °C for 5 min. Eluates and whole-cell lysates were then run a western blot analysis.

#### 3.9 Cell imaging

For exogenous foci image, HEK293T cells expressing BCL6_eGFP_ were seeded into a 6-well plate, after treated for indicated compounds and times, changed the medium with PBS, cells were imaged with Invitrogen EVOS FLoid. For immunofluorescence, 1.5 × 10^6^ Farage cells were seeded into a 6-well plate and treated with indicated compounds and times. The cells were harvested and centrifuged at 1500 rpm for 3 min, then transferred to 35 mm confocal dishes which were pre-treated with Poly-D-lysine. After 30 min, washed the un-adherent cells with PBS, fixed with 4% paraformaldehyde for 15 min and permeabilized with 0.2% Triton X100 for 15 min. Epitopes were blocked with 5% BSA for 1 h. Primary antibodies were added and incubated overnight at 4 °C. After removal of the primary antibodies and washes, secondary antibodies were added and incubated at room temperature for 60 min. Finally, the cells were stained with DAPI (Beyotime, China) for 5 min. Cells were imaged with the ZEISS LSM 900 confocal microscope.

#### 3.10 Negative stain electron microscopy (EM) analysis

To prepare grids for negative stain EM analysis of BCL6, 7.5 μM BCL6^FL^ (CUSABIO, China) in buffer (10mM Tris pH 8.0) was incubated with DMSO or 12.5 μM **D3**/**D5**/BI3802 for 1 h at room temperature. The incubated protein samples were rapidly diluted to 0.1 mg/mL (sample treated with DMSO) or 0.2 mg/ml (samples treated with **D3**/**D5**/BI-3802). A 3 μL aliquot was applied onto the glow-discharged (40 s) 300-mesh copper grid coated with a continuous carbon film (Quantifoil Micro Tools GmbH, Germany). After incubating for 1 min, the grid was blotted from sideways with a piece of filter paper and washed with 5 µL of storage buffer before staining with 0.75% uranyl formate (UF) for 40 s. The grid was positioned on a side-entry holder and loaded into a JEM-1400 operated at 120 kV, condenser lens aperture 150 μm and spot size 1. The images were taken using RADIUS software on a Morada G3 CCD camera at a magnification of 120,000 × (corresponding to a calibrated sampling of 3.23 Å per physical pixel).

#### 3.11 Knockdown of SIAH1 in HEK293T cells

HEK293T cells expressing BCL6_eGFP_ were seeded into a 6-well plate, the next day, 3.75 ul lipo 3000 transfection reagent (Invitrogen, USA) was added to 125 ul OPTI-MEM (Invitrogen, USA) (mix A). 5ul SIAH1 si-RNA (GenePharma, China) were added to 125 ul OPTI-MEM (Invitrogen, USA) (mix B). Adding mix B to mix A and incubated for 15 min at RT then added it to HEK293T cells in a dropwise manner. Cells were next cultured for 48 h. siRNAs targeting SIAH1 were designed and synthesized by Suzhou GenePharma. SIAH1 siRNA Sense: 5’-CUGUGAGUUUAGGCCUUAUTT-3’, Anti-sense: 5’-AUAAGGCCUAAACUCACAGTT-3’.

#### 3.12 Overexpression of BCL6_eGFP VSP>GSA_ in HEK293T cells

6×10^5^ HEK293T cells were seeded into a 6-well plate, the next day, 3.75 μl lipo 3000 transfection reagent (Invitrogen, USA) was added to 125 μl OPTI-MEM (Invitrogen, USA) (mix A). 5 μl BCL6_eGFP VSP>GSA_ plasmid (GenePharma, China) and 5 ul p3000 were added to 125 μl OPTI-MEM (Invitrogen, USA) (mix B). Adding mix B to mix A and incubated for 15 min at RT then added it to HEK293T cells in a dropwise manner. Cells were next cultured for 24 h.

#### 3.13 MTT

Farage cells were seeded (2×10^4^ cells per well in 100 μL of medium) in 96-well plate, then added 100 μl medium containing various concentrations of compounds to each well. After three days, 20 μl of a 5 mg/mL MTT solution was added to each well and cells were incubated for additional 2~4 h at 37℃. 50 μl of 20% (w/v) SDS was directly added to each well and cells were incubated at 37℃ overnight, the absorbance of each well was measured at 570 nm wavelength, calculate cell viability through Graphpad Prism 8.0.

#### 3.14 Edu staining

1.5×10^5^ Farage cells were seeded in 12-well plates, the medium contained 1 or 5 μM D3/BI3802. After 72 hours, the cells were incubated with the medium contained 10 µM Edu for next 2 h. Then cells were harvested and centrifuged at 1500 rpm for 3 min, then transferred to 35 mm confocal dishes which were pre-treated with Poly-D-lysine. After 30 min, washed the un-adherent cells with PBS, then stained with BeyoClick™ EdU Cell Proliferation Kit (Beyotime, China). After staining, the samples were photographed and analyzed by the ImageJ software.

#### 3.13 Cell proliferation analysis

Farage cells were seeded in 12-well plates (5 × 10^3^ cells per well) and treated with 1/5 μM **D3**/BI3802 for 9 days. The cells were counted on day 1, 3, 5, 7, and 9.

### 4. NMR and HRMS spectra of compounds

**^1^H NMR spectrum of D1**

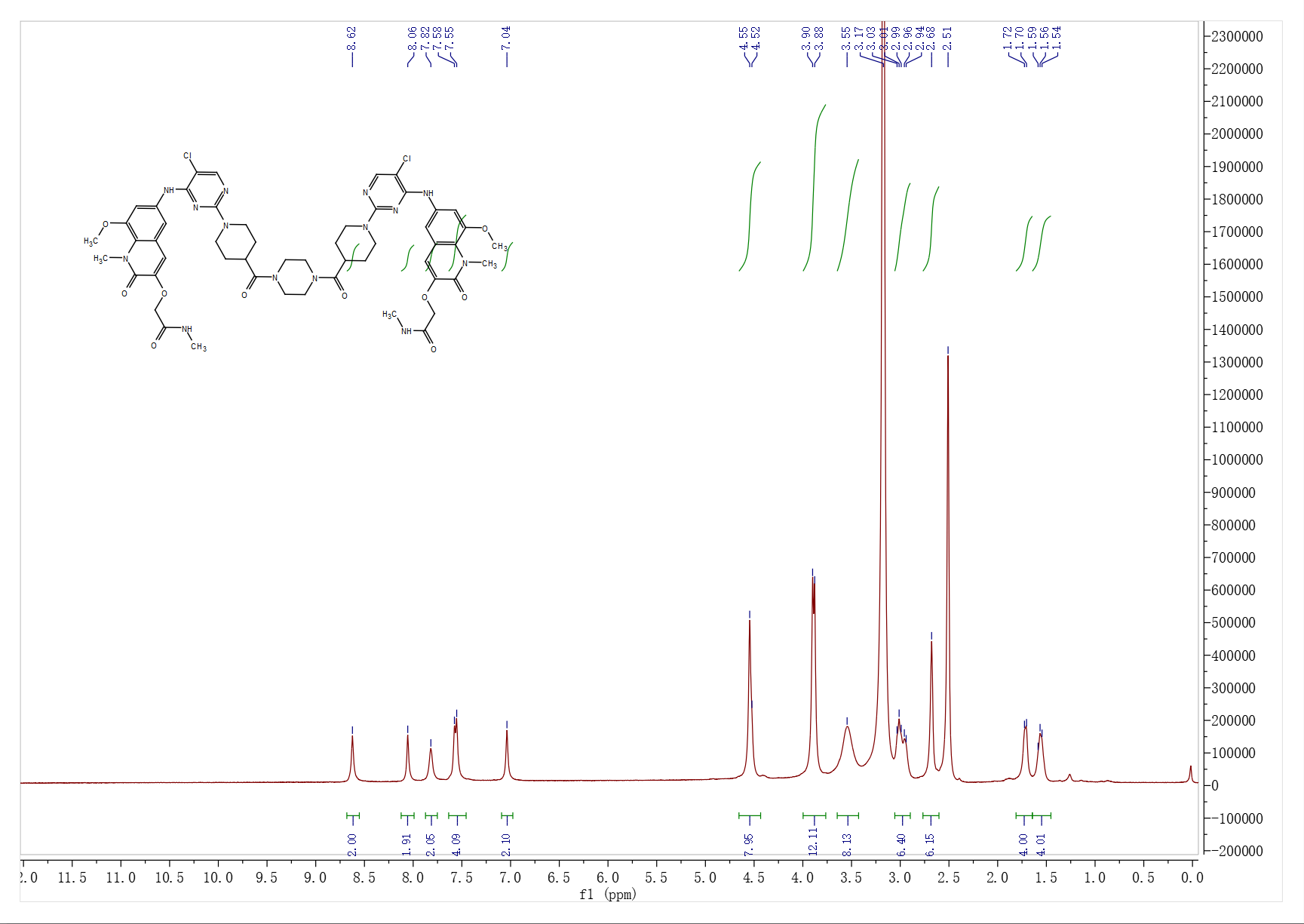

**^13^C NMR spectrum of D1**

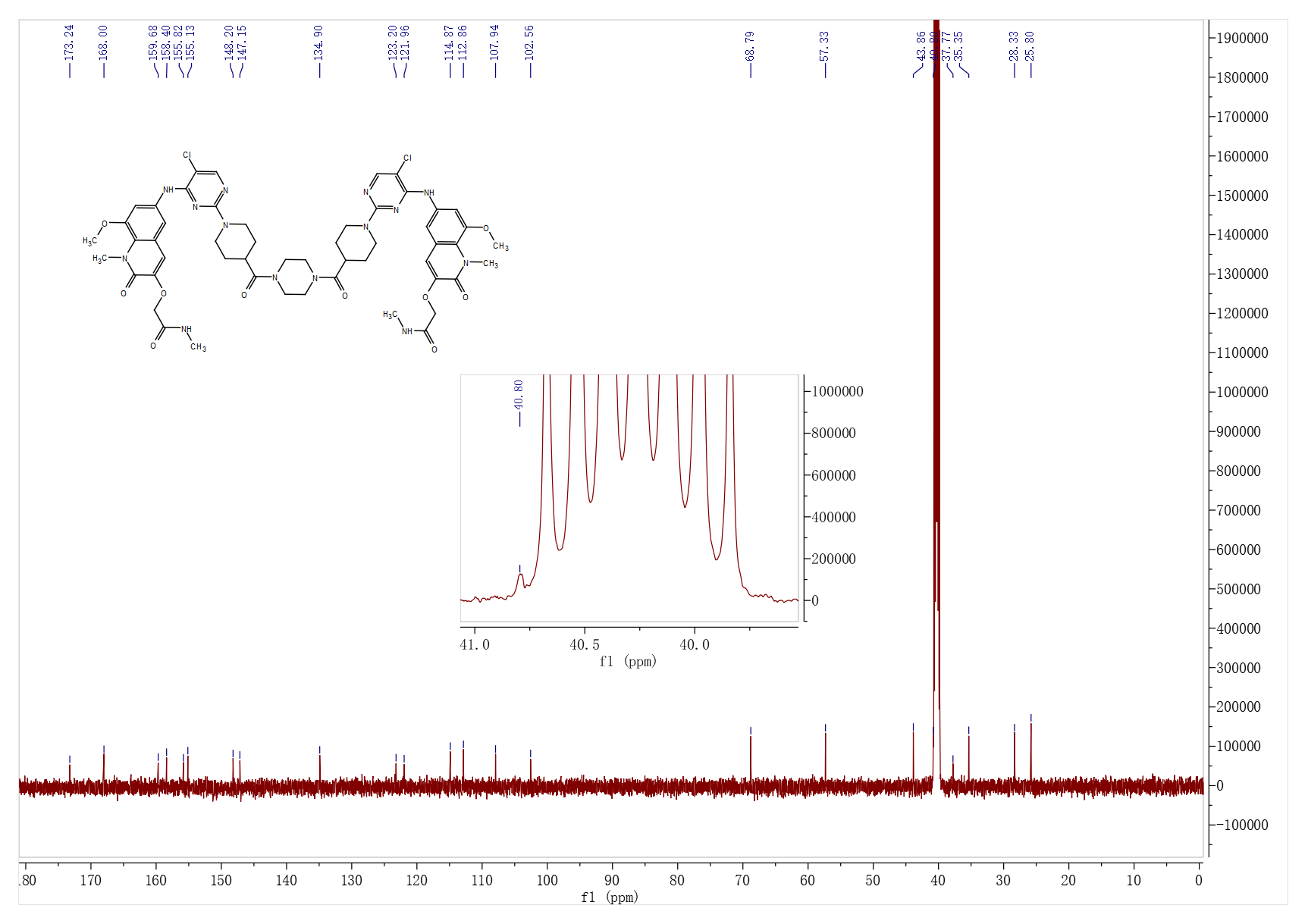

**HRMS of D1**

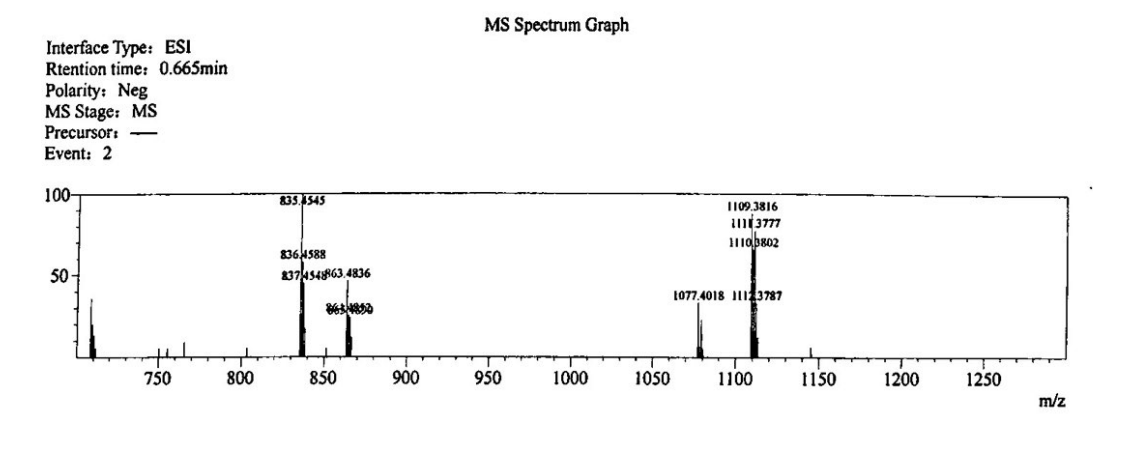

**^1^H NMR spectrum of D2**

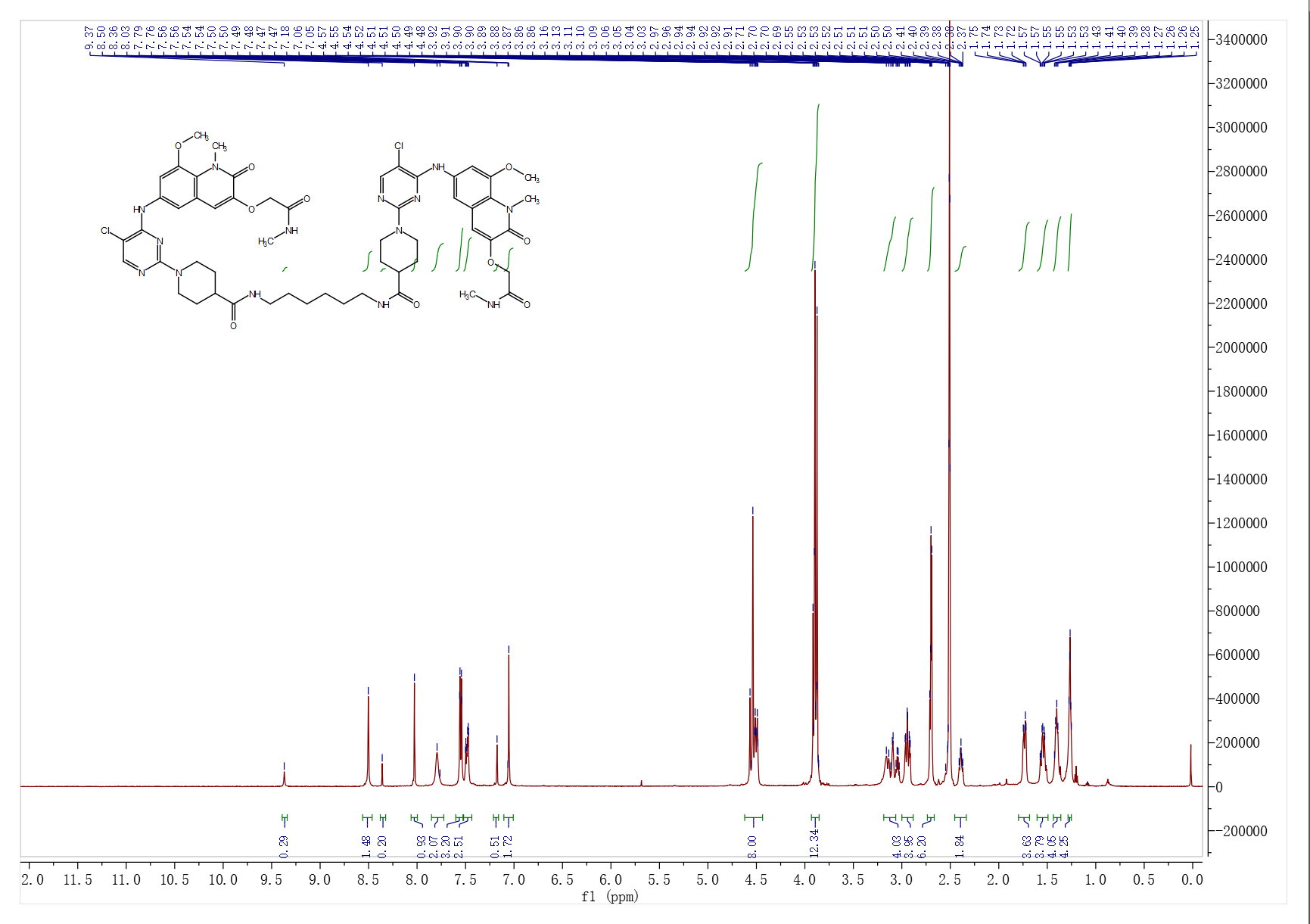

**^13^C NMR spectrum of D2**

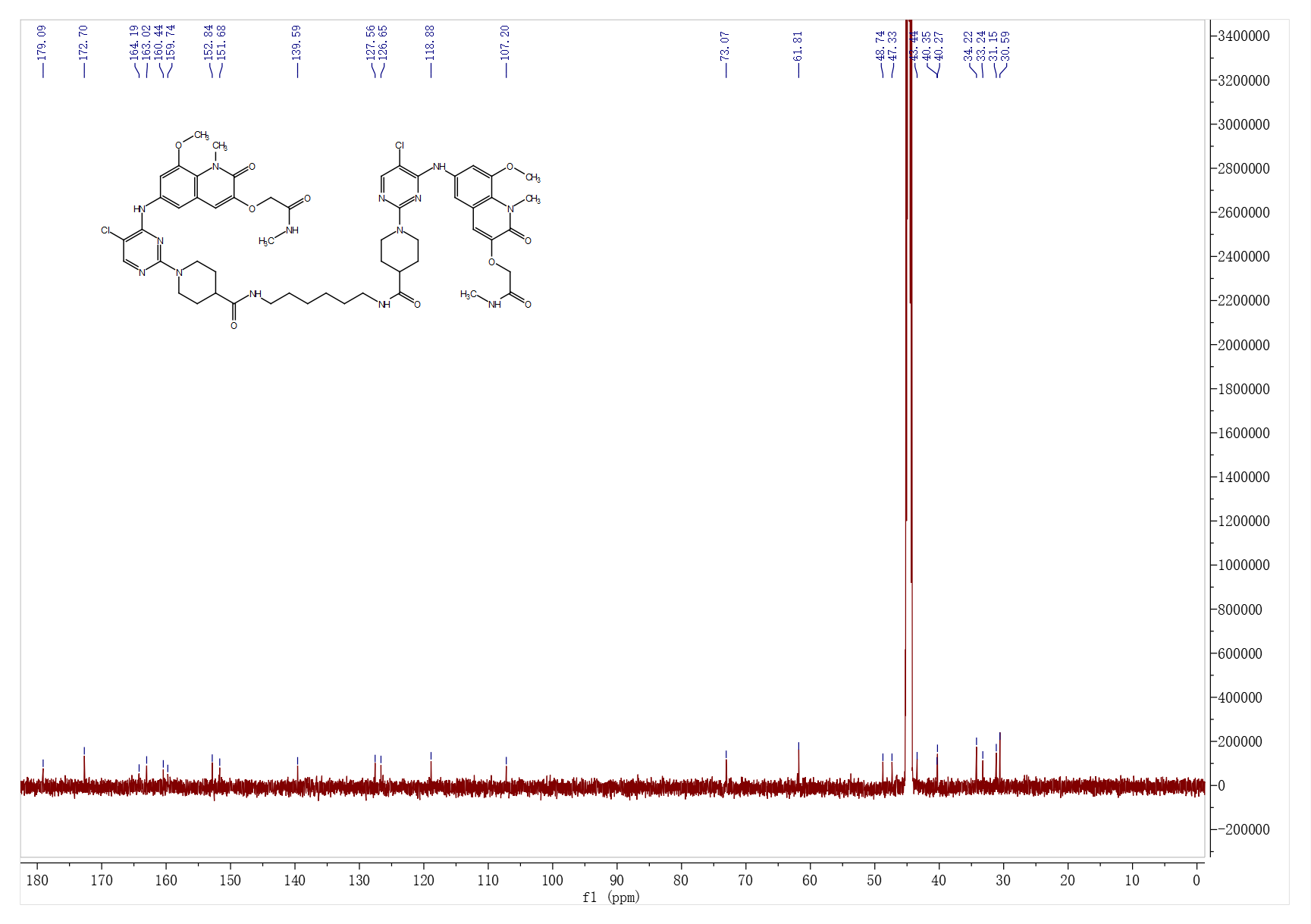

**HRMS of D2**

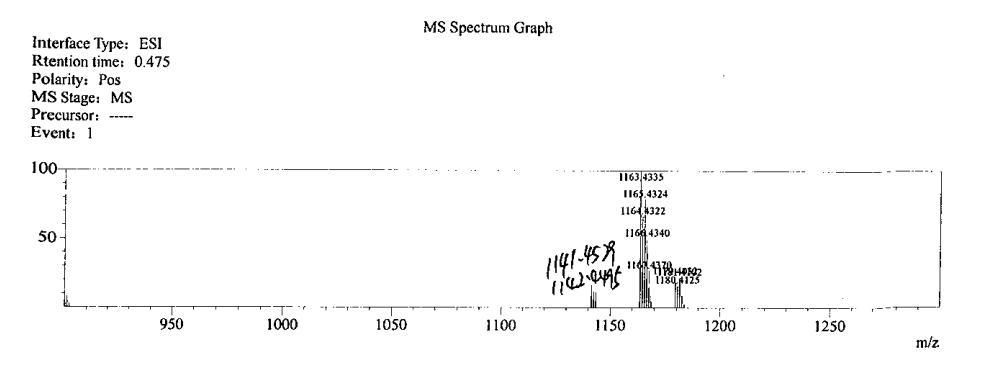

**^1^H NMR spectrum of D3**

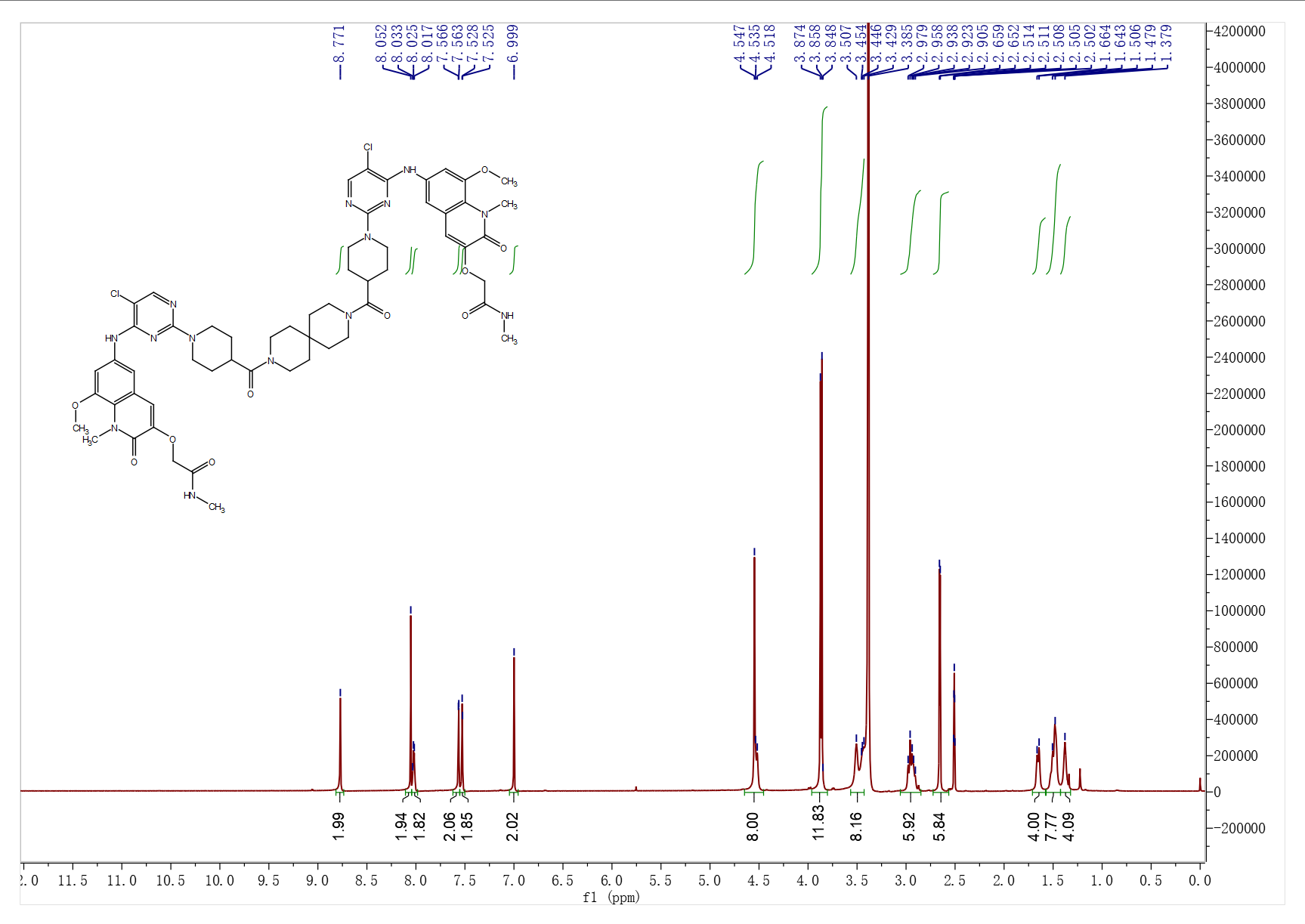

**^13^C NMR spectrum of D3**

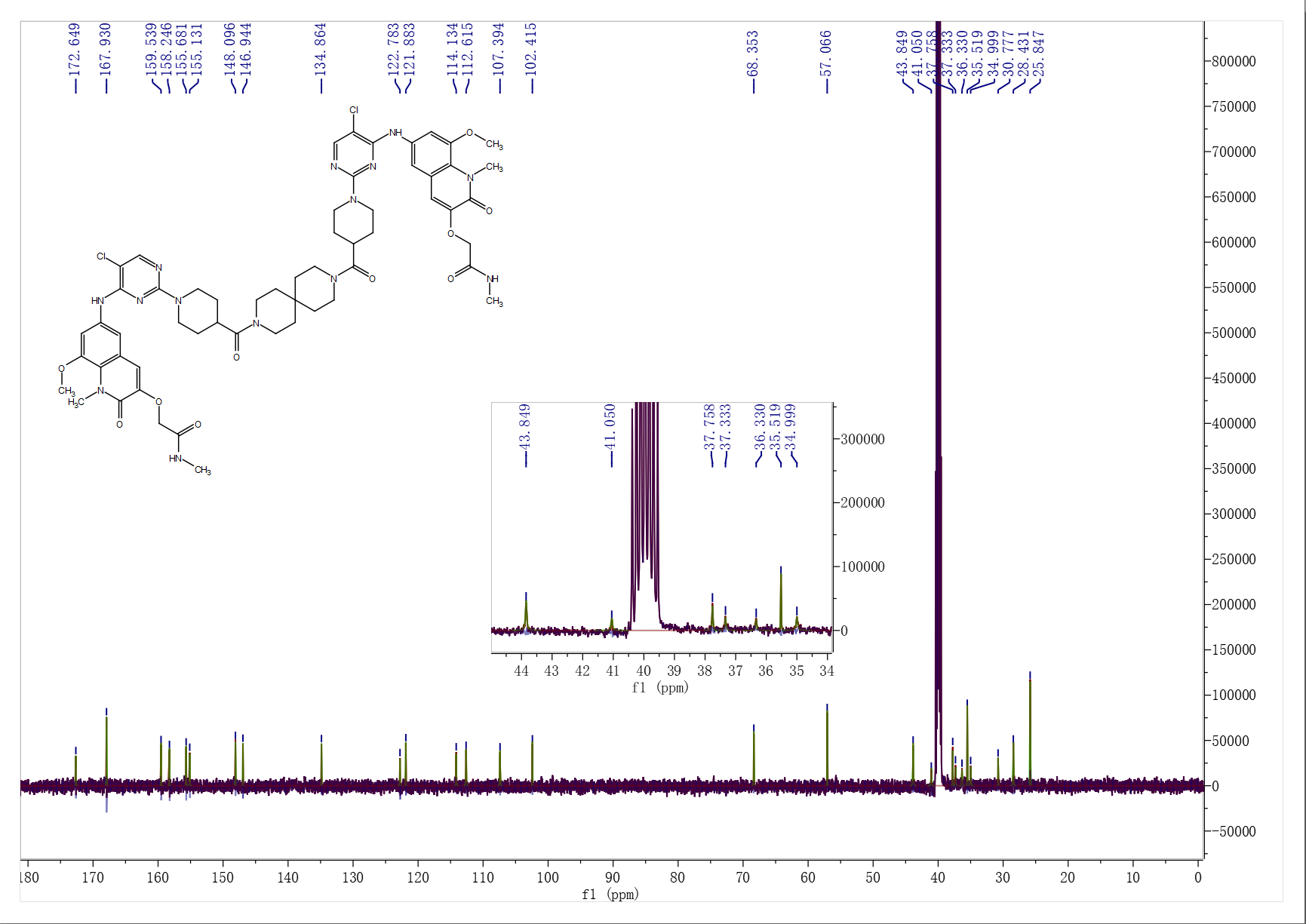

**HRMS of D3**

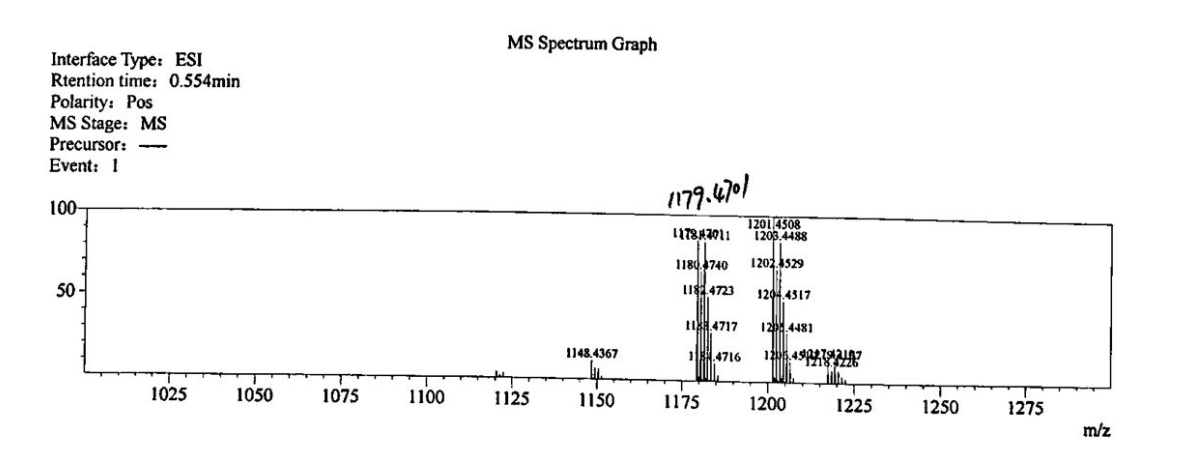

**^1^H NMR spectrum of D4**

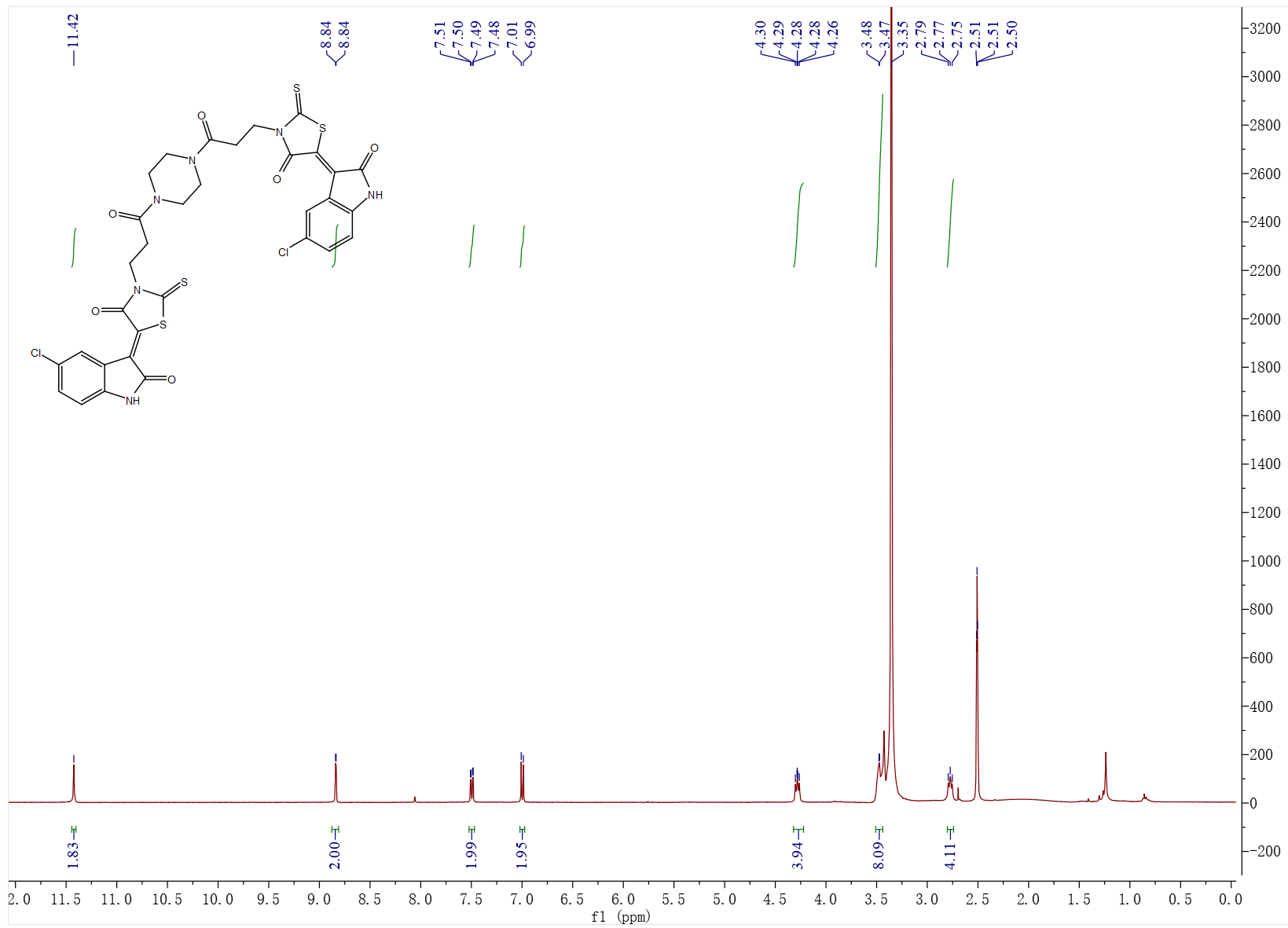

**^13^C NMR spectrum of D4**

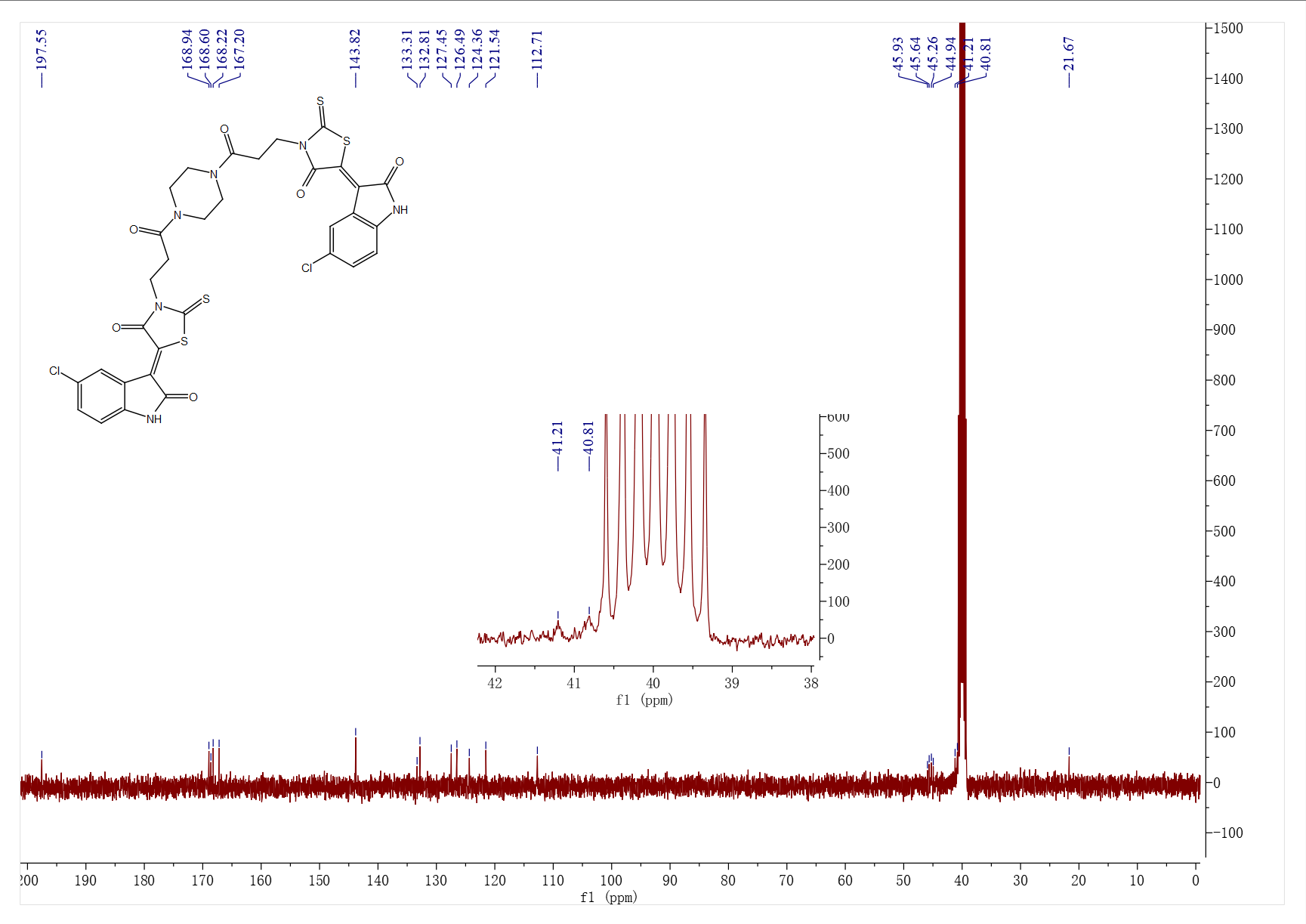

**HRMS of D4**

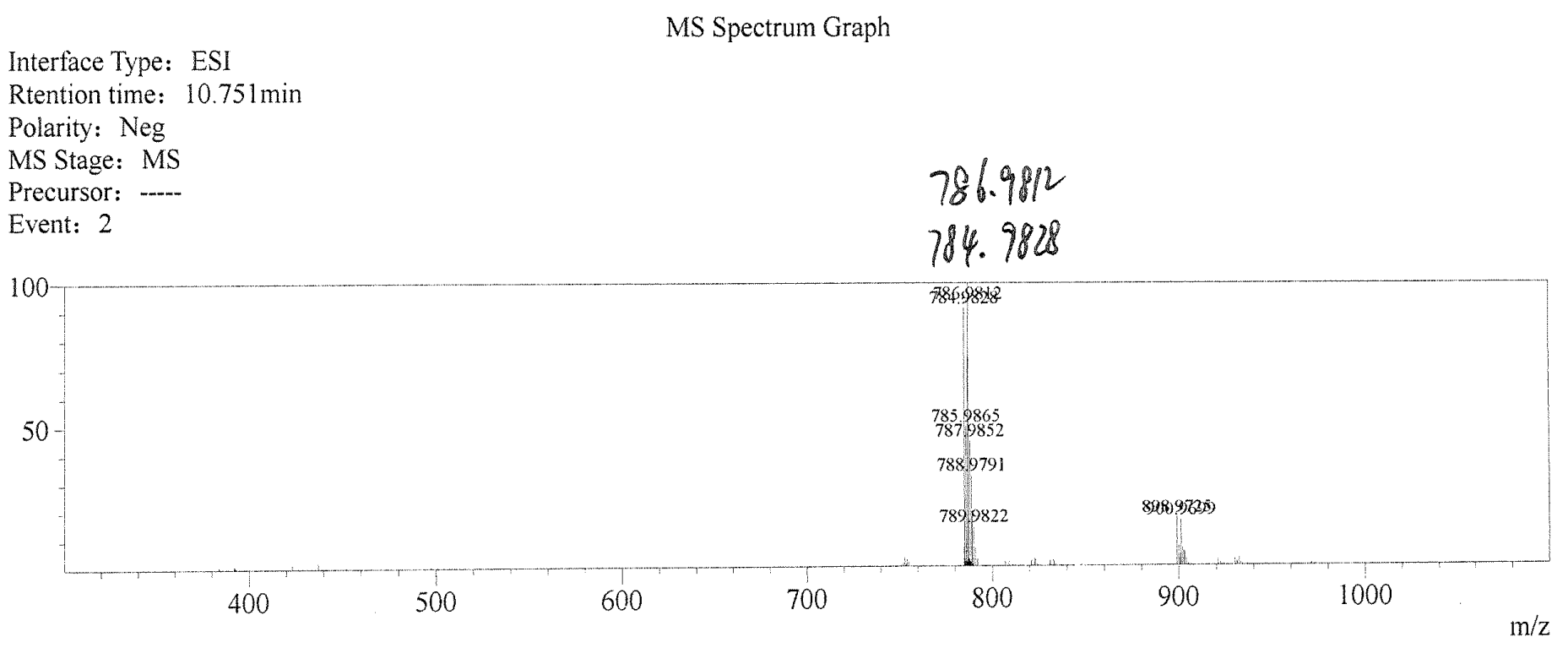

**^1^H NMR spectrum of D5**

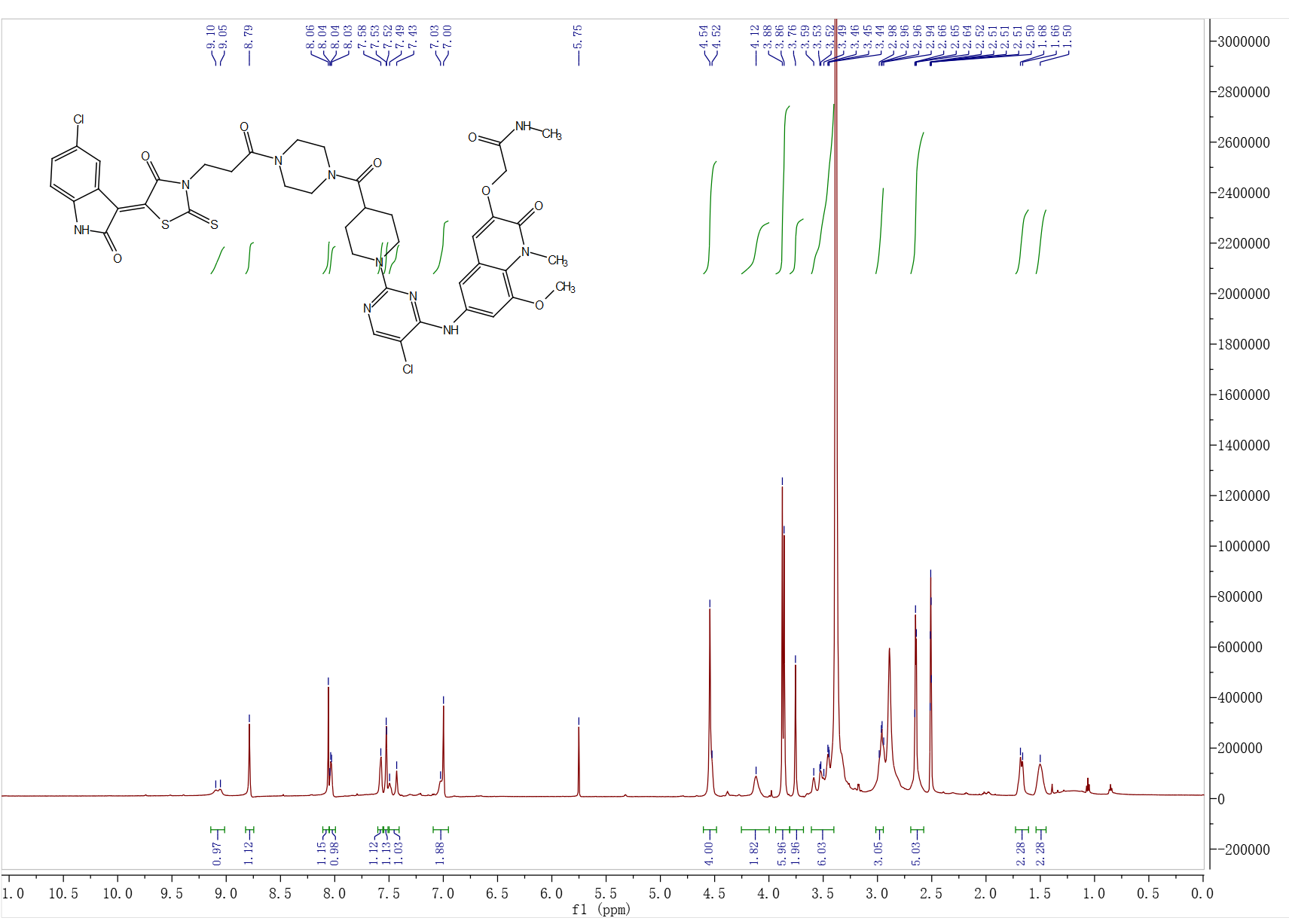

**^13^C NMR spectrum of D5**

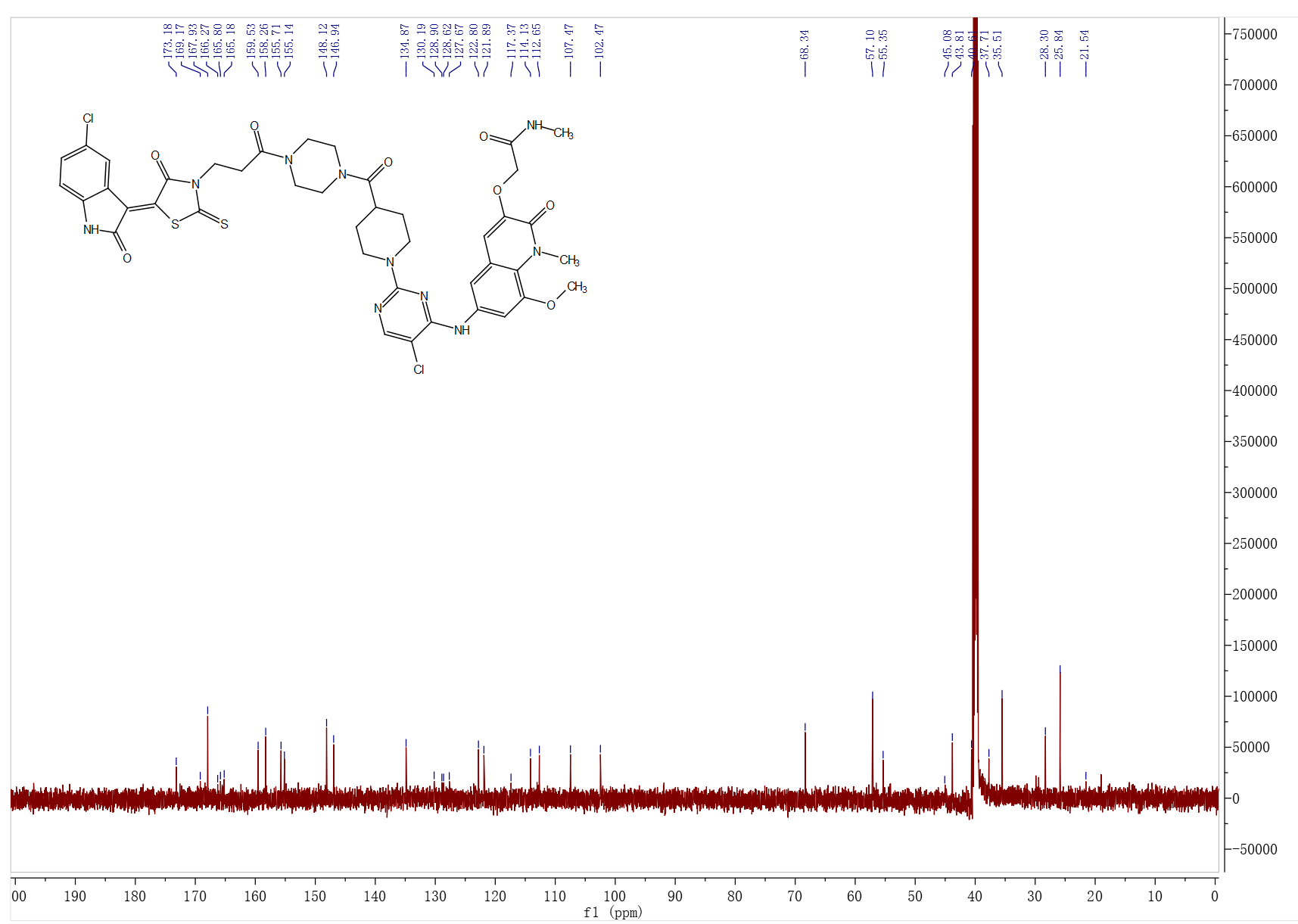

**HRMS of D5**

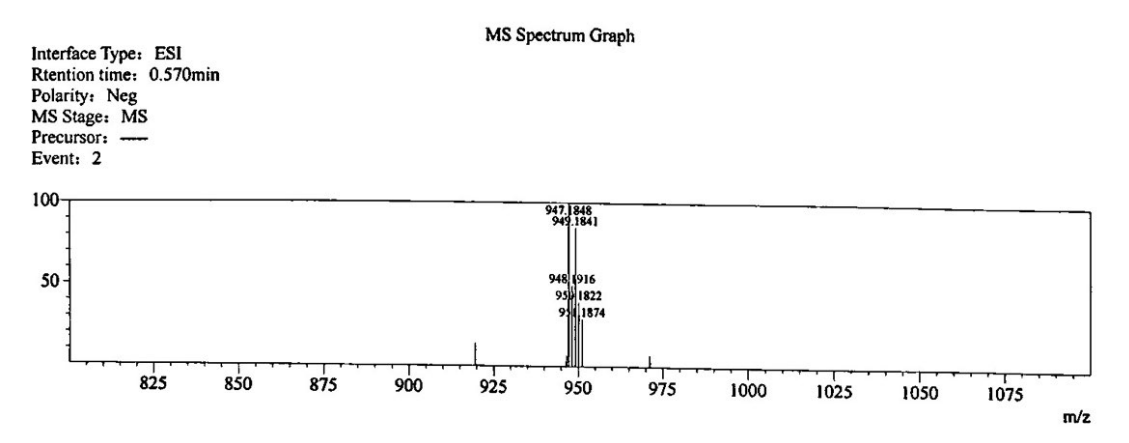
